# VAPB and its binding partner AKAP11 promote lipid droplet degradation

**DOI:** 10.64898/2026.01.27.699657

**Authors:** Shalom Borst Pauwels, Menno Spits, Lennert L.J. Janssen, Sarah Rotman, Arnoud H. de Ru, Anja W.M. de Jong, Erik Bos, Peter A. van Veelen, Roman I. Koning, Martin Giera, Jacques Neefjes, Birol Cabukusta

## Abstract

The endoplasmic reticulum (ER) is the master regulator of various cellular processes. To achieve its diverse functions, the ER interacts with other organelles at membrane contact sites, regions where organelles are brought into proximity. Most ER membrane contact sites are facilitated by the ER-resident VAP proteins. To address the role of VAP proteins in regulating cellular lipid homeostasis, we performed a targeted lipidomic screen after silencing individual VAPs. The loss of VAPB increases cellular levels of neutral lipids stored in lipid droplets (LDs). The increase in neutral lipids is reflected in the size, number and motility of LDs, and is due to the impaired degradation of these organelles. VAPB requires its contact site forming ability to regulate LDs, prompting the identification of protein kinase A (PKA) anchor AKAP11 as a VAPB interaction partner in regulating LD degradation and dynamics. Collectively, our findings identify a role for the ER-resident VAPB-AKAP11 interaction and PKA activity in regulating LD homeostasis.

**Summary:** The ER-resident membrane contact site protein VAPB and its interaction partner AKAP11 regulate lipid droplet size and motility by mediating neutral lipid degradation. VAPB requires its ability to form membrane contact sites and interact with AKAP11, protein kinase A anchor protein, for mediating lipid droplet homeostasis. This paper uncovers a role for VAPB-AKAP11-PKA axis in regulating the homeostasis of lipid droplets.

## Introduction

The endoplasmic reticulum (ER) is the largest membrane-bound organelle, extending as a network throughout the cytoplasm. The ER supports a multitude of functions including calcium signaling, organelle fission and lipid synthesis and transport (Voeltz et al., 2024; Wu et al., 2018). Although organelles are physically separated, the ER requires direct interaction with other organelles to perform these roles. These interactions occur at membrane contact sites, regions where organelles are in close proximity without fusing.

The majority of ER contact sites are facilitated by the vesicle-associated membrane protein (VAMP)-associated protein (VAP) family. The five VAP proteins – VAPA, VAPB, MOSPD1, MOSPD2 and MOSPD3 – form membrane contact sites by recruiting other proteins, and their associated organelles, to the ER (Fig. 1A). They achieve this by interacting with short linear motifs found in other proteins, so-called FFAT (two phenylalanines [FF] in an acidic tract) motifs and their related FFAT-like and FFNT (two phenylalanines in a neutral tract) motifs (Cabukusta et al., 2020; Neefjes and Cabukusta, 2021). For example, at ER-*trans* Golgi contact sites, VAPA interacts with OSBP for the non-vesicular trafficking of cholesterol to the trans Golgi, which is critical for the cholesterol pools abundant on the plasma membrane (Mesmin et al., 2013).

**Figure 1.**
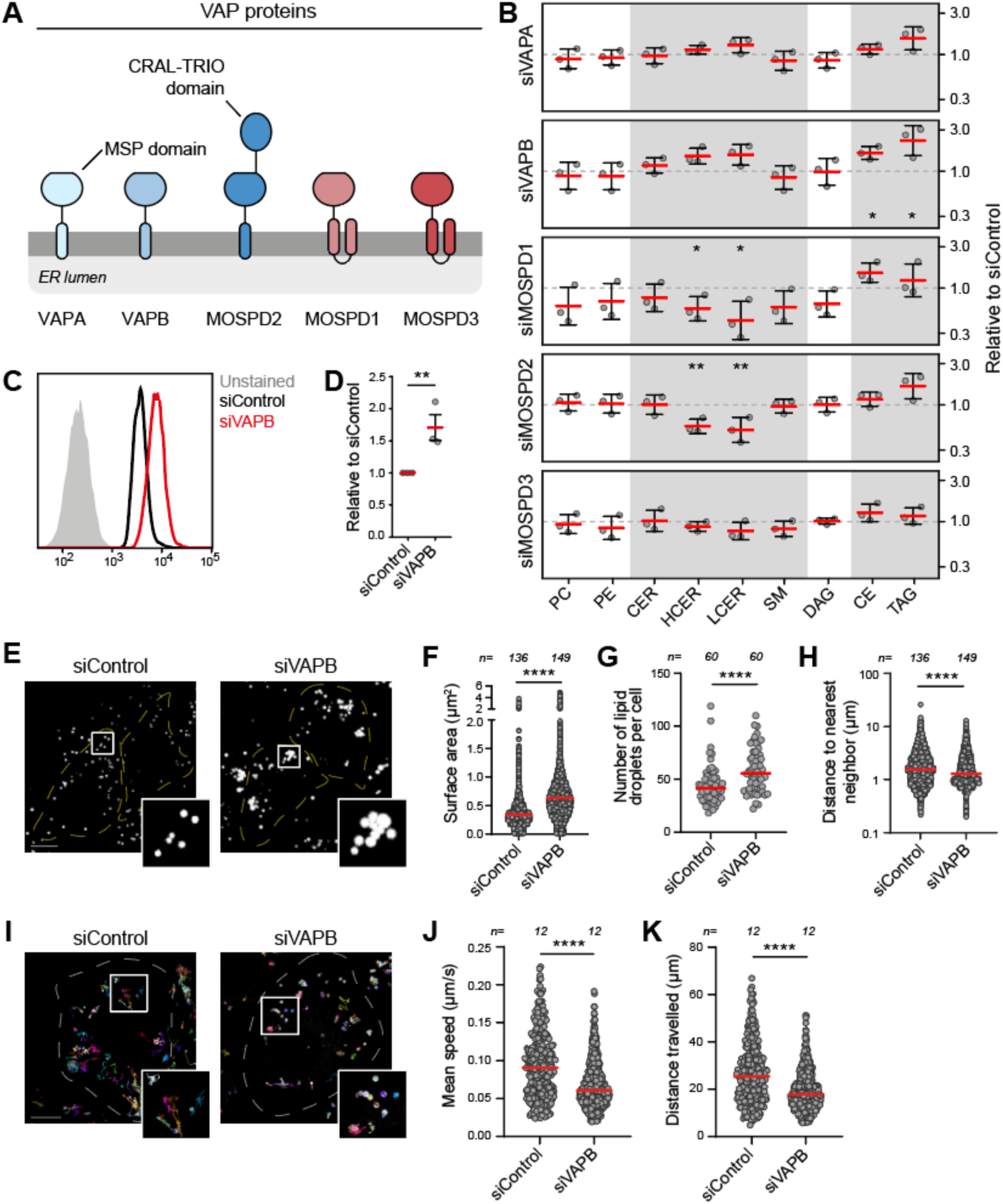
VAPB depletion increases LD size and reduces LD motility. **(A)** Schematic overview of the VAP protein family. **(B)** Lipidomic analysis after siRNA depletion of VAP proteins in HeLa cells, relative to siControl levels (Suppl. Table 1). Results are mean ± SD of three independent experiments. PC: phosphatidylcholine, PE: phosphatidylethanolamine, CER: ceramide, HCER: hexosylceramide, LCER: lactosylceramide, SM: sphingomyelin, DAG: diacylglycerol, CE: cholesterol ester, TAG: triacylglycerol. **(C)** Flow cytometry analysis of Nile Red-stained HeLa cells after VAPB silencing. **(D)** Quantifications corresponding to panel C. Mean fluorescence intensity relative to siControl is shown. Results are mean ± SD of three independent experiments. **(E)** Representative fluorescence images of siControl- and siVAPB-treated cells stained with BODIPY to visualize LDs. Scale bar: 10 µm. **(F-H)** Quantification of LD surface area (F), number of LDs per cell (G), and clustering of LDs (H) corresponding to images of panel E. Red bars represent the median of three (F and H) or four (G) independent experiments; n is the number of analyzed cells. **(I)** TrackMate tracks of LD movement for BODIPY-stained siControl-and siVAPB-treated cells imaged for 5 minutes (Suppl. Movie 1). Scale bar: 10 µm. **(J and K)** Quantification of mean speed of LDs (J), and distance travelled by LDs in 5 minutes (K) corresponding to panel I. Red bars represent the median of two independent experiments; n is the number of analyzed cells.

Lipid droplets (LDs) are ER-derived organelles specialized in storing neutral lipids (Mathiowetz and Olzmann, 2024; Olzmann and Carvalho, 2019). Their lipid-laden hydrophobic core is surrounded by a phospholipid monolayer coated with integral and peripheral proteins. LDs supply lipids to other organelles for energy production and for the production of new membranes, and they function as buffers by sequestering toxic lipids. They are highly dynamic organelles that can alternate between periods of growth and breakdown depending on the metabolic state of the cell. The genetic and metabolic dysfunction of lipid droplets is implicated in cancer, lipodystrophy and metabolic and neurodegenerative diseases (Zadoorian et al., 2023).

Here, we performed a targeted lipidomic screen to study the role of VAP proteins in lipid homeostasis. This revealed that VAPB is involved in the regulation of CE and TAG levels, and accordingly in LD dynamics. Loss of VAPB impairs the degradation of LDs, resulting in enlarged and less motile LDs, in which its ability to interact with FFAT motifs plays a critical role. By mapping the FFAT motif-targeted VAPB interactome, we identify the PKA anchor AKAP11 as a VAPB-binding partner that is likewise necessary for LD degradation. Collectively, we propose a role of VAPB-AKAP11 interaction and PKA activation in maintaining LD homeostasis.

## Results

### VAPB regulates neutral lipid levels and LD dynamics

To investigate the role of individual VAP proteins in maintaining cellular lipid homeostasis, we performed lipidomic analysis of HeLa cells, a human cervical carcinoma cell line, after siRNA depletion. The depletion of individual VAP proteins resulted in various lipid imbalances (Fig. 1B) (Suppl. Table 1). Among these, we found that MOSPD1 and MOSPD2 depletion decreased hexosylceramide and lactosylceramide levels. Moreover, levels of triacylglycerols (TAG) and cholesterol esters (CE) were significantly increased after the depletion of VAPB. This increase was found across all TAG and CE species (Fig. S1A-C). Accordingly, the increased neutral lipid levels in VAPB-depleted cells were confirmed by Nile Red staining (Fig. 1C, D).

As TAG and CE are abundantly found in lipid droplets (LDs), we investigated whether LDs are affected by VAPB loss. The depletion of VAPB increased the size of LDs, observed by the neutral lipid stain BODIPY and by immunostaining using a perilipin-2 (PLIN2) antibody, an LD coat membrane protein (Fig. 1E, F) (Fig. S2A, B). The enlarged LD phenotype was also observed in other cell lines, Hep-G2, Huh-7, MelJuSo and U2OS, suggesting a ubiquitous effect of VAPB depletion on LD size (Fig. S3A-H). The possibility of an off-target effect of the pooled siRNAs was excluded by the observation that individual VAPB-targeting siRNAs resulted in the same phenotype (Fig. S2C, D).

In addition to the enlarged LD size, we observed an increased number of LDs after VAPB depletion, further in line with the observed increase in neutral lipid levels (Fig. 1G). An increase in both LD size and number implies that the LD enlargement was not due to aberrant fusion of LDs, but due to a defect in neutral lipid metabolism. Furthermore, we observed an increased clustering of LDs in VAPB-depleted conditions, suggesting a mobility defect (Fig. 1E, H). Next, we examined the effect of VAPB depletion on LD motility. Loss of VAPB impaired LD motility compared to the control, represented by a decreased speed and travelled distance (Fig. 1I-K) (Suppl. Movie 1). Collectively, these data propose a role for the ER protein VAPB in LD metabolism.

### VAPB coordinates LD breakdown

Increased levels of neutral lipids can be a result of elevated lipid synthesis or decreased lipid breakdown. To pinpoint whether VAPB depletion affects the synthesis or degradation of neutral lipids, we performed metabolic chasing experiments using a fluorescent diacylglycerol analog (NBD-DAG). Cells were incubated with NBD-DAG for 2 hours, and its conversion to TAG (NBD-TAG) was analyzed by thin-layer chromatography (Fig. 2A) (Fig. S4A). In line with our earlier observations (Fig. 1B), VAPB depletion resulted in an accumulation of newly synthesized TAG (Fig. 2B, C). Pharmacological inhibition of LD degradation (depicted as ATGLi+LALi) negated the effect of VAPB depletion on TAG accumulation (Fig. 2B, C), (Fig. S4B). This observation suggests that neutral lipid degradation is impaired by the absence of VAPB.

**Figure 2.**
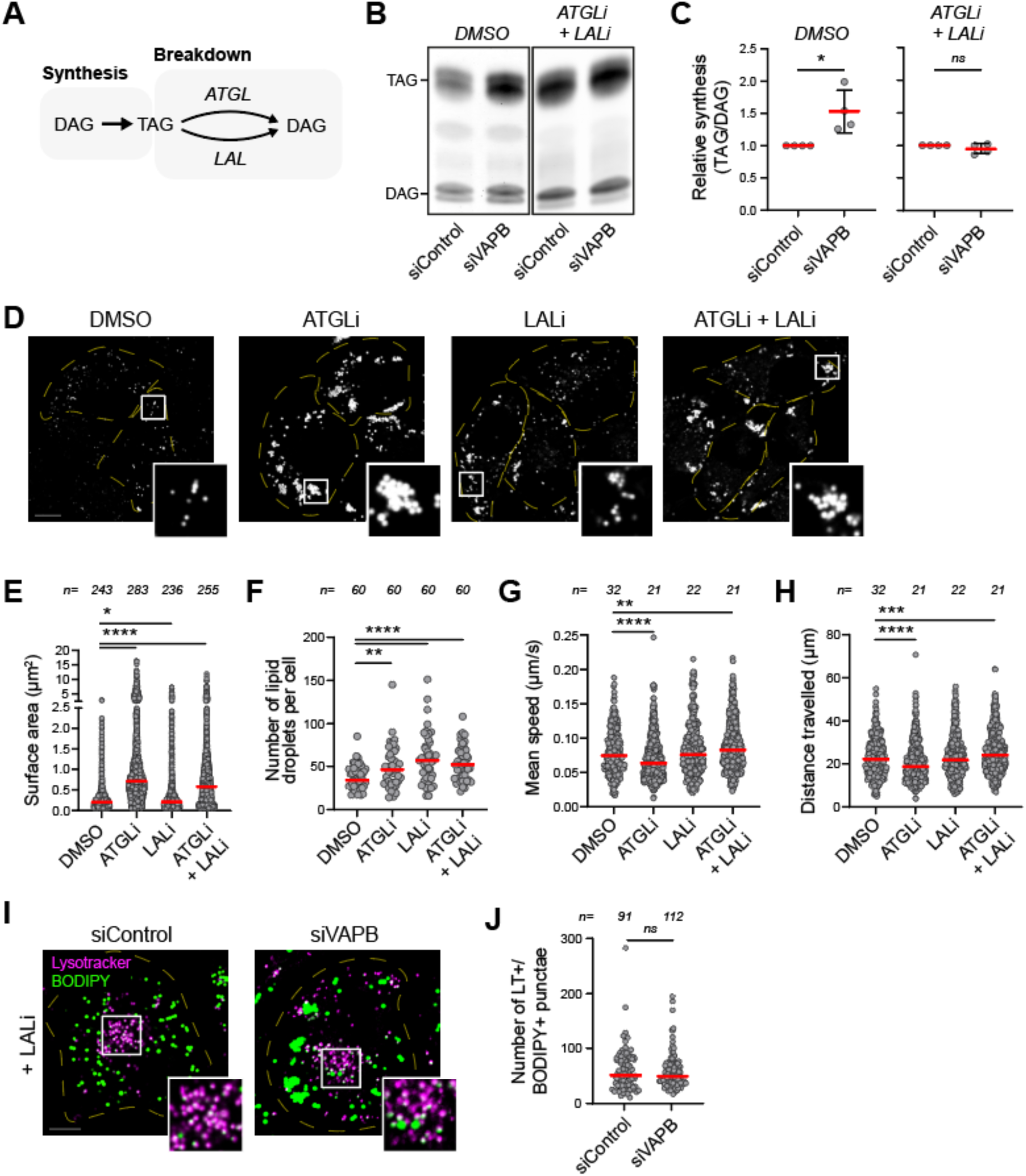
VAPB depletion impairs LD degradation. **(A)** Schematic representation of TAG metabolism. TAG is synthesized from DAG and can be degraded back to DAG by ATGL and LAL. **(B)** Representative thin-layer chromatography image of metabolic chasing experiments. siControl- and siVAPB-treated cells were fed with NBD-DAG for 2 hours in the presence or absence of ATGLi and LALi. Lipids were extracted, separated on a thin-layer chromatography plate, and visualized using fluorescence. **(C)** Quantification of NBD-TAG relative to NBD-DAG corresponding to images of panel B. Red bars represent the mean ± SD of three independent experiments. **(D)** Representative fluorescence images of cells treated with ATGLi or LALi, stained with BODIPY to visualize LDs. Scale bar: 10 µm. **(E-H)** Quantification of LD surface area (E), number of LDs per cell (F), mean speed of LDs (G), and distance travelled by LDs in 5 minutes (H) (Suppl. Movie 2). Red bars represent the median of three independent experiments; n is the number of analyzed cells. **(I)** Fluorescence images of cells incubated with LALi for 16 hours and stained with BODIPY and lysotracker to label LDs and lysosomes, respectively. Scale bar: 10 µm. **(J)** Number of puncta double positive for lysotracker (LT) and BODIPY corresponding to images of panel I. Red bars represent the mean of three independent experiments; n is the number of analyzed cells.

LD degradation reportedly occurs through two pathways: lipolysis mediated by adipose triglyceride lipase (ATGL) and lipophagy—the direct uptake of LDs by lysosomes (microlipophagy) and the uptake of LDs by autophagosomes, later fusing with lysosomes (macrolipophagy) (Mathiowetz and Olzmann, 2024). In lysosomes, LDs are degraded by lysosomal acid lipase (LAL). We wondered which degradation pathway is affected by VAPB depletion. We inhibited ATGL or LAL and observed their effect on LDs. ATGL inhibition resulted in enlarged LDs and an increased number of LDs, similar to VAPB-depleted cells (Fig. 2D-F). While inhibition of LAL or the combined inhibition of ATGL and LAL (referred to as ATGL+LAL) also led to an increased number of LDs, LDs did not show an increase in size to the same extent as in ATGL inhibition. Additionally, ATGL inhibition decreased motility of LDs, while LAL and ATGL+LAL inhibitions did not (Fig. 2G, H) (Suppl. Movie 2). Collectively, ATGL inhibition resulted in a similar LD phenotype observed in VAPB depletion, suggesting that VAPB is involved in ATGL-mediated LD degradation. We found that VAPB depletion does not affect ATGL localization to LDs, nor ATGL protein levels, suggesting a role of VAPB in an alternative regulatory mechanism (Fig. S5A and B).

LALi treatment renders lysosomes unable to degrade LDs, resulting in the accumulation of neutral lipids within lysosomes (Fig. S6). Supporting the notion that VAPB depletion affects ATGL-mediated LD degradation, the number of BODIPY-positive lysosomes following LAL inhibition was not affected by VAPB depletion, implying that VAPB is not involved in the uptake of LDs by lysosomes (Fig. 2I, J). Together, these data suggest that VAPB mediates LD breakdown through lipolysis.

### VAPB is a component of ER-LD contact sites

VAP proteins facilitate membrane contact sites by tethering organelles to the ER. While MOSPD2 interacts directly with LDs via its CRAL-TRIO domain, VAPA and VAPB interact with other proteins, to bridge ER and LDs (Zouiouich et al., 2022). FFAT motif-containing ORP2, VPS13A and VPS13C tether ER to LDs (Kumar et al., 2018; Wang et al., 2020; Yeshaw et al., 2019). Present at ER-LD-mitochondria tripartite contact sites, the MIGA2-VAPB complex plays a role in LD synthesis (Freyre et al., 2019).

To confirm the association of the ER-resident protein VAPB with LDs, we tagged the *VAPB* gene with mScarlet in its endogenous locus in HeLa cells (Fig. 3A) (Fig. S2E). Live-cell imaging of endogenous VAPB with LDs revealed instances in which VAPB-enriched ER subdomains were colocalizing with LDs, indicating the presence of VAPB at ER-LD contact sites (Fig. 3B). We then questioned whether VAPB is necessary for contact between ER and LDs. Electron microscopy images of control- and VAPB-depleted cells showed that neither the percentage of LDs in contact with the ER, nor the length of ER-LD contact sites was affected by VAPB depletion (Fig. 3C-E) (Fig. S7, 8). Although VAPB takes part in ER-LD contact sites, it is not necessary for the creation or the architecture of these sites.

**Figure 3.**
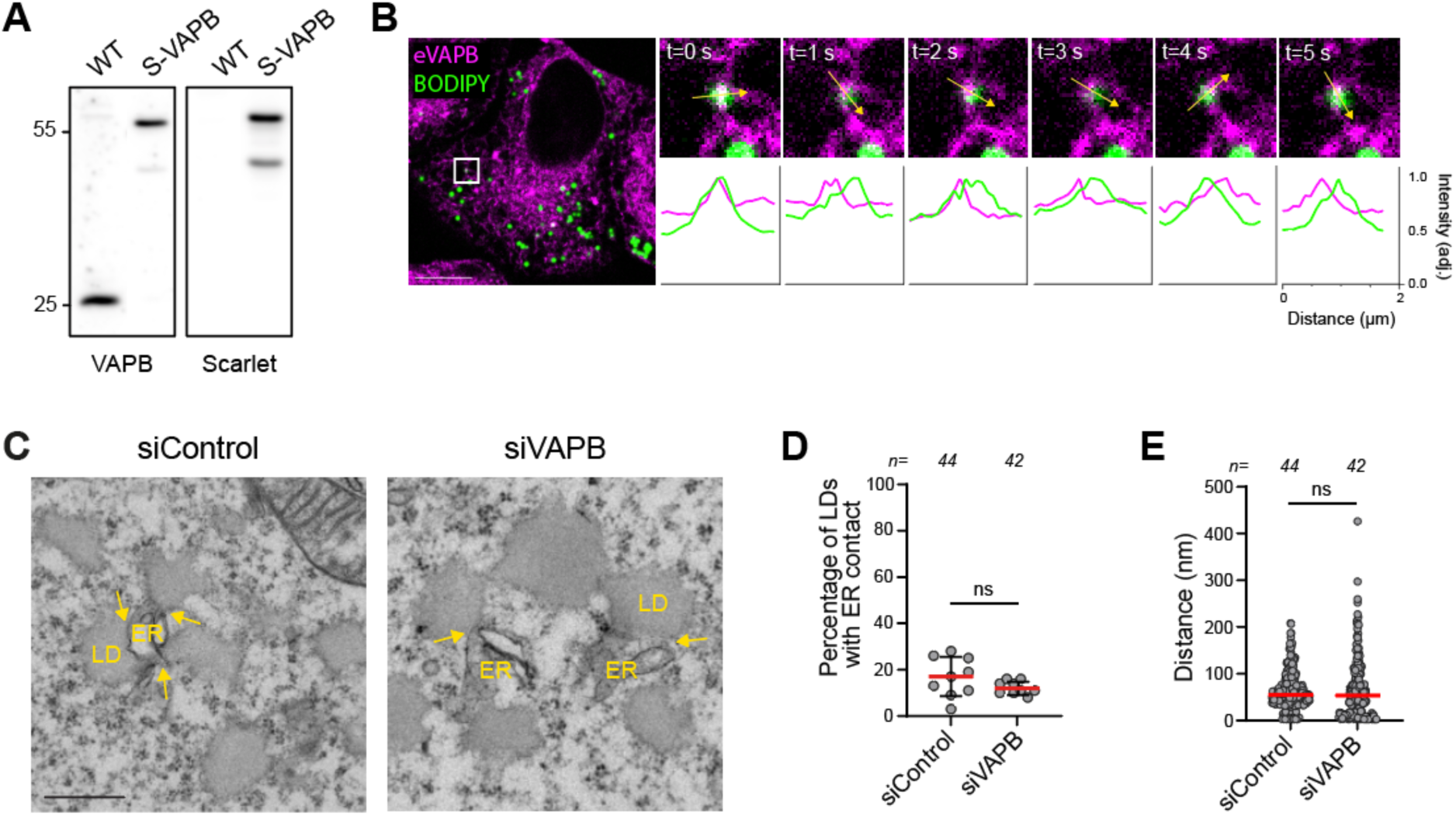
VAPB is present at ER-LD contact sites and does not affect its architecture. **(A)** Western blot validation of endogenous tagging of VAPB with mScarlet (S-VAPB). **(B)** Fluorescence images of mScarlet-VAPB cells stained with BODIPY, visualizing the contact of endogenous VAPB (eVAPB)-positive ER domains with LDs (t in seconds). Scale bar: 10 µm. **(C)** Electron microscopy images of ER-LD contact (indicated by arrows) in siControl and siVAPB cells. Scale bar: 500 nm. **(D-E)** Quantification of the percentage of LDs in contact with the ER in a given electron microscopy section (D), and the length of ER-LD contacts (Fig. S5) (E). Red bars represent the mean ± SD in panel D and the median in panel E; n is the number of analyzed cells.

### VAPB regulates LD size through FFAT motif binding

VAPB recruits binding partners to the ER by interacting with their FFAT motifs. We investigated whether the ability to interact with FFAT motifs is necessary for VAPB’s role in LD homeostasis. We performed rescue experiments, in which various RFP-labelled VAPB mutants were expressed in VAPB-depleted cells (Fig. 4A). Expression of wild-type VAPB rescued the increase in LD size, further validating our observations. Meanwhile, the VAPB^K87D, M89D^ mutant that fails to interact with FFAT motifs, depicted as VAPB**, was unable to revert the LD phenotype, showing that VAPB requires its FFAT motif binding ability to regulate LD homeostasis (Fig. 4B, C). In line with this observation, a similar result was obtained with a VAPB truncation consisting of the protein’s transmembrane region, VAPB-TM. These data show that the FFAT motif binding ability of VAPB is critical for LD homeostasis.

**Figure 4.**
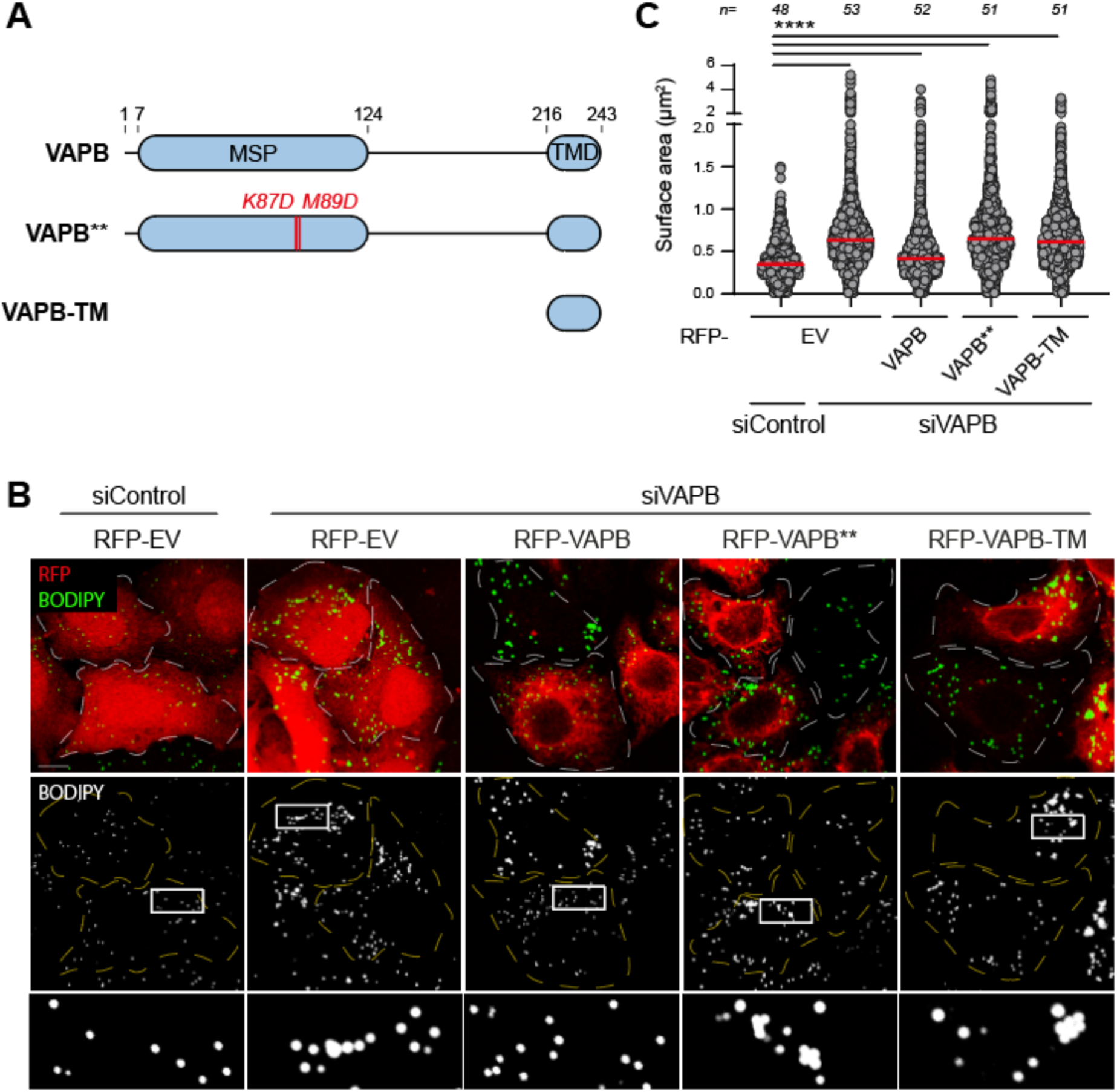
FFAT motif binding by VAPB is necessary for regulating LD size. **(A)** Domain architecture of VAPB, VAPB**: a VAPB mutant unable to interact with FFAT motifs, and VAPB-TM. MSP: major sperm protein, TMD: transmembrane domain. **(B)** Fluorescence images of siRNA-treated cells expressing RFP-empty vector (RFP-EV) or various RFP-tagged VAPB proteins. Scale bar: 10 µm. **(C)** Quantification of LD surface area from images corresponding to images of panel B. Red bars represent the median of three independent experiments; n is the number of analyzed cells.

### Motif-targeted VAPB proteome identifies VPS13A, ORP1L and AKAP11 as LD regulators

Based on the previous observation, we aimed to identify interaction partners of VAPB that are involved in regulating LD metabolism. We utilized the TurboID method that applies proximity-based biotinylation of interactors, followed by affinity purification and proteomics identification (Fig. 5A) (Branon et al., 2018). As we were interested in interaction partners that may regulate LD homeostasis, we incubated cells with oleic acid to stimulate LD formation. Additionally, we expressed VAPB** and VAPB-TM, that are unable to rescue LD size, as negative controls (Fig. S9A). Using this method, we identified proteins selectively enriched in the interactome of wild-type VAPB compared to the negative controls (Fig. 5B) (Fig. S9B, C).

**Figure 5.**
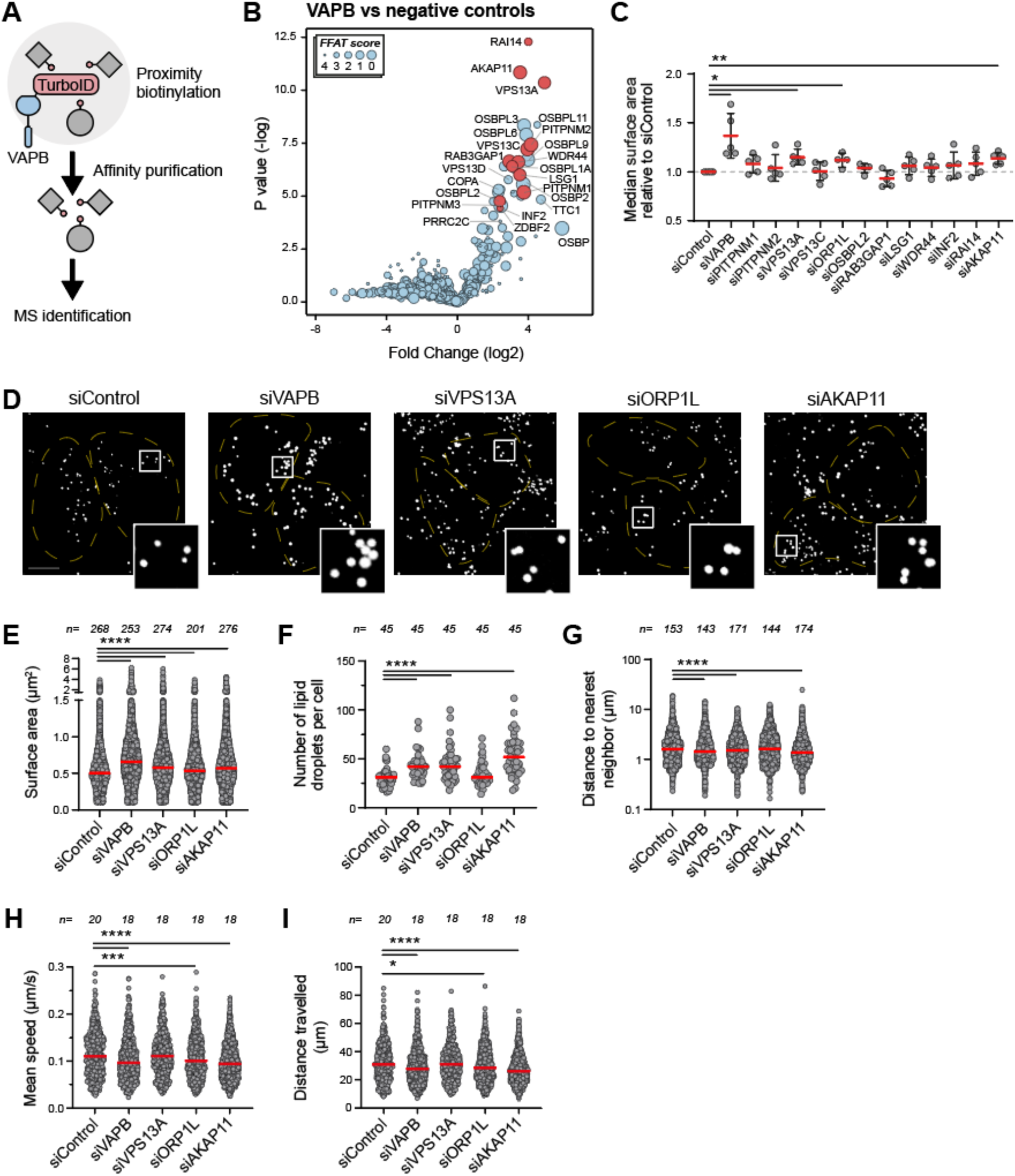
The identification of VAPB interaction partners in regulating LD dynamics. **(A)** Schematic representation of the TurboID method. TurboID cloned onto VAPB biotinylates proteins in close proximity, which are enriched by affinity purification and subsequently identified by mass spectrometry (MS). **(B)** Volcano plot of proteins enriched in wild-type VAPB in comparison to the negative controls (non-transfected, TurboID only, VAPB** and VAPB-TM). Proteins chosen for validation are depicted in red. Lower FFAT score corresponds to higher resemblance to the canonical FFAT motif. **(C)** Quantification of the median surface area of LDs in siRNA-treated cells relative to siControl. Results are mean ± SD of four or five independent experiments. **(D)** Representative fluorescence images of siControl-, siVAPB-, siVPS13A-, siORP1L- and siAKAP11-treated cells stained with BODIPY to visualize LDs. Scale bar: 10 µm. **(E)** Quantification of LD surface area. Red bars represent the median of five independent experiments (four for siORP1L); n is the number of analyzed cells. **(F-I)** Quantification of number of LDs per cell (F), and clustering (G), mean speed (H) and travelled distance in 5 minutes (I) of LDs (Suppl. Movie 3). Red bars represent the median of three independent experiments; n is the number of analyzed cells.

We selected a subset of proteins involved in a variety of cellular processes to investigate their involvement in LD size regulation (Fig. 5C) (Mikitova and Levine, 2012). Some of these proteins also contain FFAT motifs, likely facilitating an interaction with VAPB. From this selection, the depletions of VPS13A, ORP1L (*OSBPL1A*) and AKAP11 resulted in enlarged LDs, a phenotype similar to VAPB silencing (Fig. 5D, E).

VPS13A, a bridge-like lipid transfer protein, localizes to LDs, and loss of which affects LD abundance (Chen et al., 2022; Yeshaw et al., 2019). ORP1L is a lysosomal cholesterol sensor that regulates lysosomal positioning and lysosome-autophagosome fusion, but its role in LD homeostasis remains elusive (Rocha et al., 2009; Wijdeven et al., 2016). AKAP11 is a member of the AKAP family, which scaffold protein kinase A (PKA) to specific subcellular locations to spatiotemporally organize PKA signaling (Omar and Scott, 2020). Recently, loss of AKAP11 was reported to increase LD abundance in astrocytes (Liu et al., 2025).

In addition to enlarged LDs, we observed increased numbers of LDs after the silencing of VPS13A and AKAP11, and this was accompanied by increased LD clustering (Fig. 5F, G). Meanwhile, LD number and clustering were not affected by ORP1L depletion. We then examined LD motility and observed a decreased LD speed and travelled distance in ORP1L- and AKAP11-depleted cells, while these parameters were not affected in VPS13A-depleted cells (Fig. 5H, I) (Suppl. Movie 3). As VPS13A-, ORP1L- or AKAP11-depleted cells demonstrated different LD phenotypes, we concluded that these proteins regulate LD homeostasis via distinct pathways.

### VAPB coordinates LD degradation through AKAP11

VPS13A, ORP1L and AKAP11 each contain FFAT motifs, and they interact with VAPB (Fig. 6A) (Lee et al., 2025; Liu et al., 2025; Rocha et al., 2009; Song et al., 2025; Yeshaw et al., 2019). Their ability to interact with VAPB and the effect of their silencing on LD size, number and motility suggested that VAPB regulates LD degradation together with one of these proteins. To test this, we examined whether loss of VPS13A, ORP1L and AKAP11 also lead to a defect in LD degradation. We observed that AKAP11 depletion resulted in an accumulation of newly synthesized TAG, while VPS13A and ORP1L depletion did not affect TAG synthesis (Fig. 6B, C). Similar to VAPB depletion, inhibition of LD degradation negated the effect of AKAP11 depletion on TAG accumulation (Fig. 6D, E). The depletion of AKAP11 simulates VAPB silencing in all measured parameters, metabolic activity and LD phenotype by microscopy, suggesting that AKAP11 and VAPB regulate LD degradation via the same pathway.

**Figure 6.**
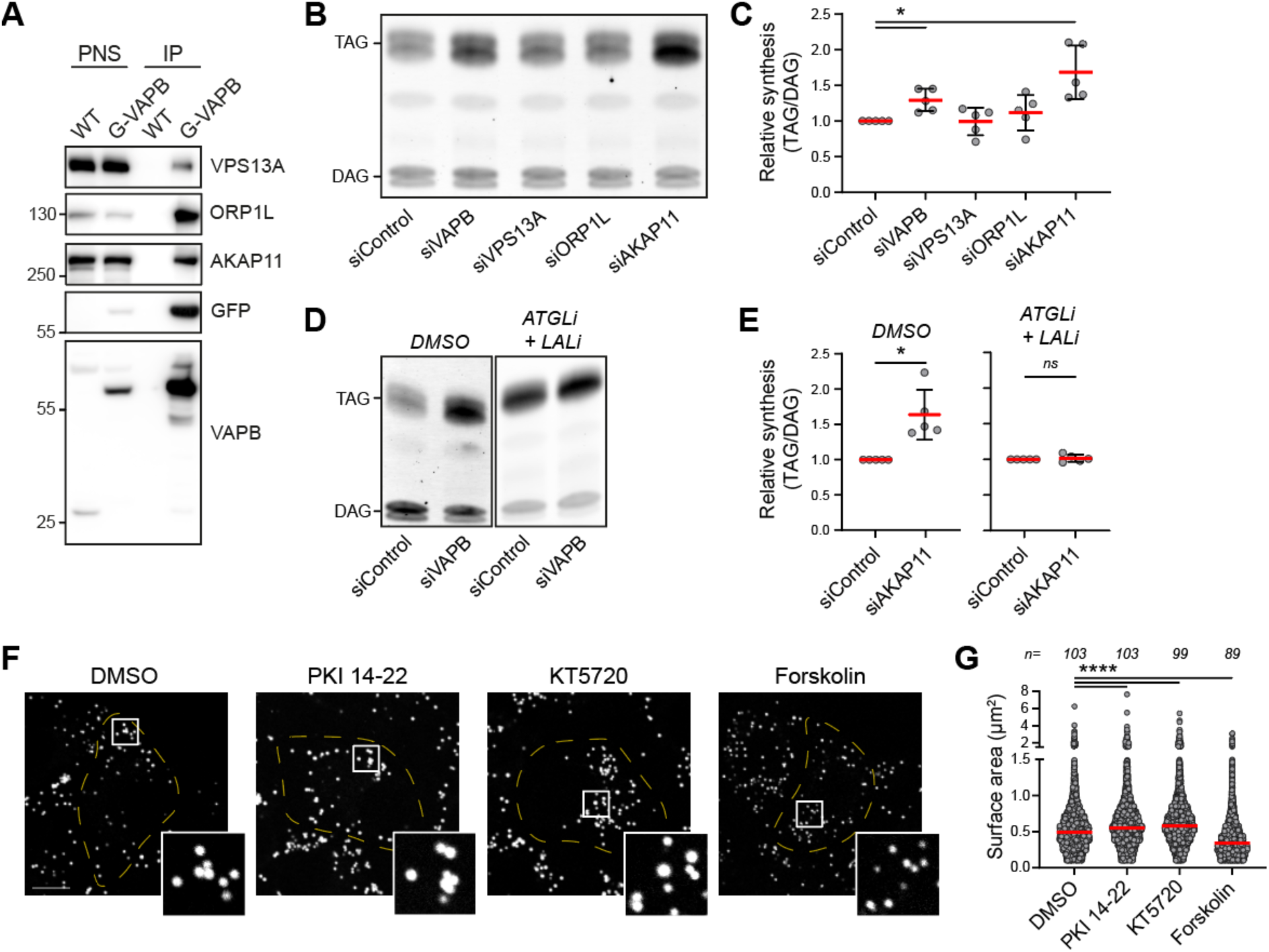
AKAP11 interacts with VAPB and is necessary for LD degradation. **(A)** Co-immunoprecipitation of endogenous GFP-VAPB (G-VAPB) and VPS13A, ORP1L and AKAP11 in MelJuSo cells. **(B)** Representative thin-layer chromatography image of metabolic chasing experiments of siRNA-treated cells fed with NBD-DAG for 2 hours. **(C)** Quantification of NBD-TAG relative to NBD-DAG corresponding to panel B. Red bars represent the mean ± SD of five independent experiments. **(D)** Representative thin-layer chromatography image of metabolic chasing experiments. siControl- and siAKAP11-treated cells were fed with NBD-DAG for 2 hours in the presence or absence of ATGLi and LALi. **(E)** Quantification of NBD-TAG relative to NBD-DAG corresponding to panel D. Red bars represent the mean ± SD of five independent experiments. **(F)** Fluorescence images of cells treated with PKI 14-22, KT 5720 and Forskolin for 24 hours, stained with BODIPY to visualize LDs. Scale bar: 10 µm. **(G)** Quantification of LD surface area corresponding to images of panel G. Red bars represent the median of three independent experiments; n is the number of analyzed cells.

AKAP11 functions as a selective autophagy receptor by interacting with LC3B and recruiting PKA for autophagic degradation (Deng et al., 2021; Zhou et al., 2024). Through scaffolding PKA and promoting its autophagic-dependent degradation, AKAP11 regulates PKA signaling (Segura-Roman et al., 2025). This led us to examine if the impaired LD degradation seen in AKAP11-depleted cells is due to aberrant PKA signaling. Cells incubated with the PKA inhibitors PKI 14-22 and KT 5720 showed enlarged LDs, while cells incubated with the PKA activator Forskolin demonstrated decreased LD size, suggesting that PKA has a regulatory function on LD size (Fig. 6F, G). Collectively, our data support a model where AKAP11 and VAPB interaction regulates LD homeostasis by mediating neutral lipid degradation through PKA signaling (Fig. 7).

**Figure 7.**
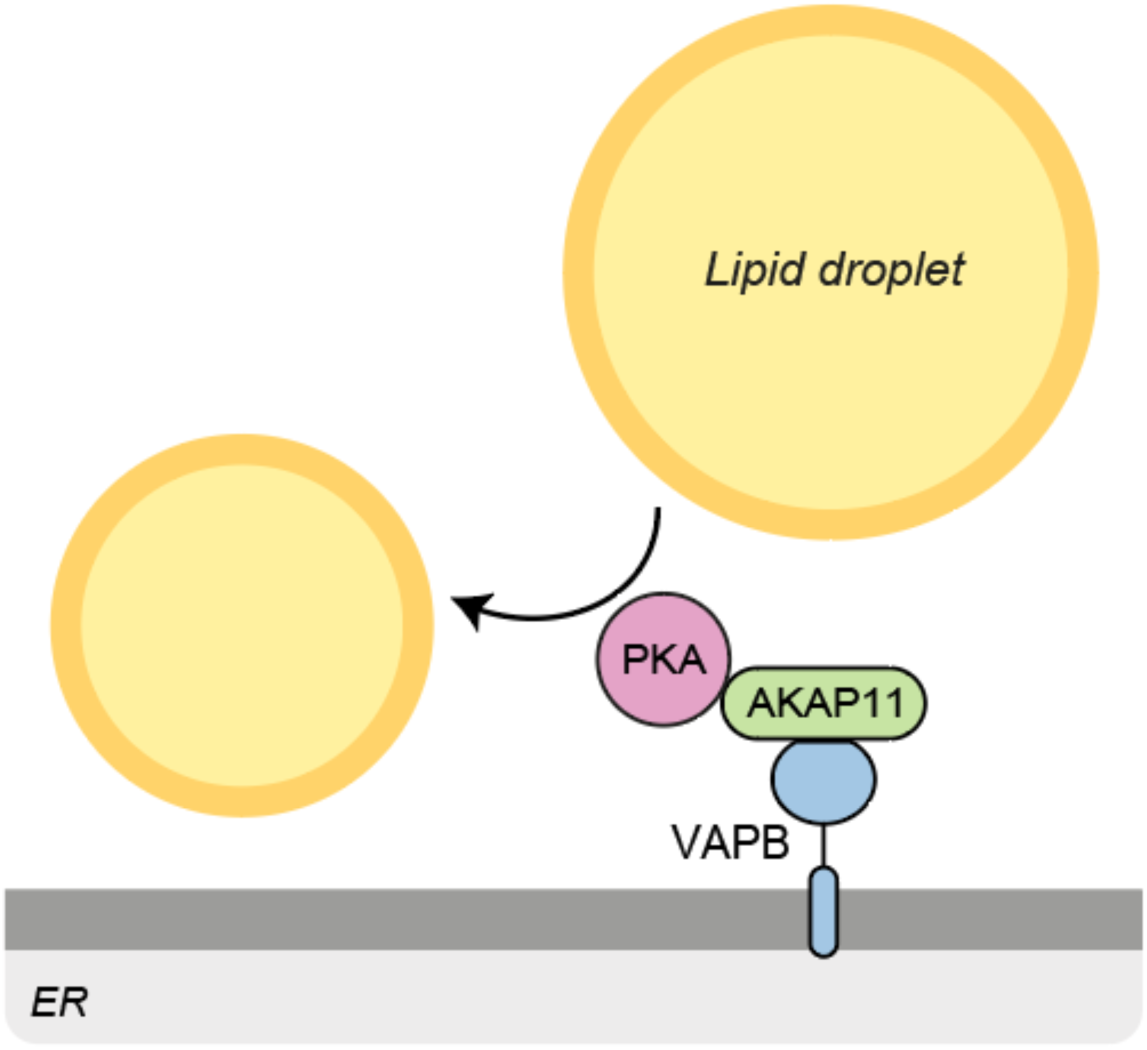
Interaction of VAPB-AKAP11 interaction promotes the degradation of LDs through PKA signaling.

## Discussion

The ER is in continuous, yet dynamic contact with other organelles to carry out various functions. The majority of these contact sites are facilitated by the ER-resident VAP proteins that interact with other organelles. In this study, we identify VAPB as a regulator of LD homeostasis that mediates their degradation; VAPB’s ability to bind FFAT motifs is essential for this function. Without VAPB, LDs are larger, more abundant and less motile. Furthermore, we identify the PKA anchor AKAP11 as a possible functional interaction partner of VAPB in mediating LD degradation through PKA activation.

AKAP11 has recently been shown to play a role in lipid metabolism, where its loss dysregulates cholesterol and fatty acid metabolic pathways and increases lipid levels, including TAG and CE (Liu et al., 2025). Loss of AKAP11 in astrocytes increases LD abundance by which excess lipid levels are sequestered into LDs as a protective mechanism against lipotoxicity. Our data show that AKAP11 plays a role in LD degradation as well. AKAP11 regulates PKA signaling by scaffolding PKA and mediating its autophagic degradation (Segura-Roman et al., 2025). The involvement of PKA in LD degradation is well established (Grabner et al., 2021; Mathiowetz and Olzmann, 2024; Schott et al., 2022). In lipolysis, PLIN1 phosphorylation dissociates its interaction partner CGI-58 (also known as ABHD5), which then interacts with ATGL for the activation of its TAG hydrolase activity. PKA also phosphorylates hormone sensitive lipase (HSL) to increase its localization on LDs and to stimulate its lipolytic activity. HSL has a broad substrate specificity including TAG, DAG, monoacylglycerols (MAG) and CE, with highest activity against DAG and CE. We show that PKA activates LD degradation, possibly following AKAP11 recruitment by VAPB.

The ER plays a central role in regulating LD biogenesis and this function is extensively studied. In contrast, its role in LD degradation and spatial organization is less well understood. Multiple proteins at ER-LD contact sites facilitate LD degradation: ESYT1 and ESYT2 remove newly hydrolyzed lipids from LDs for β-oxidation, and ORP2 mediates ATGL transport from the ER to LDs to facilitate LD degradation (Bezawork-Geleta et al., 2025; Wang et al., 2020). Additionally, VPS13D facilitates fatty acid transfer at LD-mitochondria contact sites (Wang et al., 2021). These proteins also localize to other membrane contact sites or take part in tripartite contact sites involving the ER and LDs (Giordano et al., 2013; Guillen-Samander et al., 2021; Janer et al., 2024; Wang et al., 2019). Moreover, ESYT1 and ESYT2 form a complex with VAPB, and ORP2 and VPS13D contain FFAT motifs for interacting with VAP proteins—highlighting the role of VAP proteins in LD degradation.

In addition to defected neutral lipid degradation, we found LDs are more clustered and less motile upon VAPB silencing. The link between these observations, i.e. whether they are consequential or simultaneously occurring due to loss of ER interaction, is to be elucidated. LD dispersion led by the motor proteins kinesin-1 and kinesin-2 facilitate LD contact with other organelles for degradation, which is inhibited upon microtubule disruption (Herms et al., 2015; Rai et al., 2017). Based on these observations, it is possible that loss of ER contact first leads to LD motility defects followed by impaired degradation. Indeed, the ER orchestrates the positioning and motility of various other organelles (Cabukusta and Neefjes, 2018; Jongsma et al., 2024; Lee et al., 2020; Rocha et al., 2009; Wenzel et al., 2022; Wijdeven et al., 2016). For example, the cholesterol sensor ORP1L at ER-lysosome contact sites defines lysosomal positioning, motility, and maturation. Such mechanism—an ER-bound LD sensor—could oversee LD motility and positioning prior to LD degradation.

Several mutations in *VAPB* are the cause of a familial type of the motor neuron degeneration disease amyotrophic lateral sclerosis (ALS) (Borgese et al., 2021). The T46I and P56S mutations are the two best characterized mutations and cause a dominantly inherited familial form of ALS. Neutral lipid accumulation and other lipid dysregulations are a common feature in neurodegenerative diseases, including ALS (Agrawal et al., 2022; Burg and Van Den Bosch, 2025; Feringa et al., 2025; Long et al., 2025). LDs may contribute to ALS pathogenesis through their functions in energy homeostasis, oxidative stress, and protein aggregate clearance (Farmer et al., 2020; Pennetta and Welte, 2018). We show that VAPB impacts neutral lipid levels by regulating LD degradation. Whether this mechanism is impaired in ALS pathogenesis poses an interesting research avenue. Furthermore, AKAP11 is a risk gene for schizophrenia and bipolar disorder and affected individuals show altered lipid profiles (Liu et al., 2023; Modesti et al., 2025; Palmer et al., 2022; Schneider et al., 2017; Zorkina et al., 2024). Dysregulation of LD homeostasis is suggested to play a significant role in bipolar disease pathophysiology (Pereira et al., 2024). We and others found that AKAP11 regulates LD homeostasis, indicating that impaired LD degradation could contribute to AKAP11-associated schizophrenia and bipolar disorder, possibly providing an interesting target for therapeutic manipulation (Liu et al., 2025).

## Methods

### Cell culture and transfection

HeLa, U2OS, Hep-G2 and Huh-7 cells were cultured in DMEM (Gibco #41966) supplemented with 8% fetal calf serum (Serana #S-FBS-CO-015). MelJuSo cells were cultured in IMDM (Gibco #21980) supplemented with fetal calf serum. All cell lines were authenticated by STR analysis. All experiments were performed in HeLa cells unless stated otherwise.

For siRNA transfections, serum-free medium containing DharmaFECT 1 Transfection Reagent was added to 0.5 μM siRNA and incubated for 30 minutes, constant shaking, at room temperature. Cells were added to the mixture to bring the final siRNA concentration to 50 nM. Samples were processed after 72 hours.

For plasmid transfections, cells were transfected using X-treme GENE HP DNA Transfection Reagent. Solutions containing transfection reagent and plasmid DNA were mixed, vortexed and incubated for 20 minutes at room temperature before adding dropwise to cells.

### Lipidomic analysis

HeLa cells were transfected with siRNAs and cultured in DMEM supplemented with delipidated serum (Sigma #F7524). Quantitative lipid analysis was carried out using comprehensive, quantitative flow-injection based shotgun lipidomics, employing differential mobility spectroscopy as described in (Cabukusta et al., 2024). Briefly, cells were pelleted and mixed with deuterated internal standards in methyl-tert-butyl-ether (MTBE). Lipids were extracted by an MTBE-based phase separation method. Lipids in the organic layer were analyzed using flow-injection based quantitative lipidomics. Using the Shotgun Lipidomics Assistant (SLA) software, individual lipid concentrations were calculated after correction for their respective internal standards.

### Nile Red staining and FACS analysis

HeLa cells were trypsinized and subsequently stained with 10 μg/mL Nile Red. Samples were analyzed using a BD LSR-II and data was analyzed using FlowJo software.

### Immunofluorescence staining and imaging

For fixed-cell imaging, cells were cultured on glass coverslips and fixed with 4% PFA in PBS (w/v) and quenched with 25 mM Tris-HCl pH 8.0. For LD staining, cells were permeabilized with permeabilization buffer (0.1% TritonX-100 in PBS) for 45 minutes and subsequently stained with 1μg/mL BODIPY in PBS. Samples were washed thrice with PBS and mounted onto glass slides using VECTASHIELD Vibrance Antifade Mounting Medium (Vector Laboratories #H-1700). For antibody staining, cells were permeabilized with permeabilization buffer for 30 minutes, and incubated with primary antibody diluted in permeabilization buffer for 1 hour. Cells were washed thrice with permeabilization buffer and were incubated with secondary antibody for 1 hour. After washing, LDs were stained and coverslips were mounted onto glass slides. For PLIN2 staining, cells were fixed in ice-cold methanol for 15 minutes. Images were acquired using a confocal Zeiss LSM 900 with Airyscan microscope and were analyzed and quantified using ImageJ software.

For live-cell imaging, cells were cultured in glass bottom dishes. Cells were stained with BODIPY to visualize LDs and with LysoTracker Deep Red to visualize lysosomes. Images were acquired using an Andor Dragonfly 200 or an Andor Dragonfly 500 microscope with a climate chamber at 37°C and 5% CO_2_. For observing LD motility, images were acquired every second for 5 minutes. Images were analyzed and quantified using ImageJ software.

### Analysis of LD phenotype

Cells were transfected with siRNAs or treated with 20 μM NG-497 (ATGLi) or 50 μM Lalistat-2 (LALi) for 72 hours or 10 μM PKI 14-22, 1 μM KT5720 or 10 μM Forskolin for 24 hours. MelJuSo cells were incubated with 30 μM oleic acid to stimulate LD biogenesis. LD size was quantified using the MRI Lipid Droplets Tool ImageJ plugin. Alternatively, Ilastik was employed to annotate LDs after which a mask was created in ImageJ. Watershed was applied and Analyze Particles function was used to calculate the surface area of the LDs. The Nearest Neighbor Distance ImageJ plugin was used to quantify LD clustering. As individual LDs were indistinguishable within tightly clustered LDs, LD size may reflect clustered LDs and the nearest neighbor distance may be an underrepresentation of actual LD clustering. TrackMate was used to quantify LD motility in ImageJ.

### Lysosomal uptake of LDs

Cells were incubated with 50μM Lalistat-2 (LALi) for 16 hours. The lysosomal uptake of LDs was analyzed by quantifying the number of BODIPY and LysoTracker double positive puncta per cell. For analysis, masks were generated for BODIPY and LysoTracker images. The masks were overlayed using Image Calculator and quantified using Analyze Particle function in ImageJ.

### Metabolic chasing of NBD-DAG and thin-layer chromatography

HeLa cells were transfected with siRNAs and cultured in a 24-well plate. Cells were washed with PBS and incubated with serum-free DMEM containing inhibitors for 30 minutes: 20 μM ATGLi and 50 μM LALi or 20 μM DGAT1i and 20 μM DGAT2i. Cells were then incubated with the inhibitors, 30 μM oleic acid and 10 μM NBD-DAG for 2 hours. After incubation, cells were washed and trypsinized, to which 4% NaCl was added. Lipids were extracted via the Bligh and Dyer method as described in (Cabukusta et al., 2024).

After lipid extraction, dried lipids were resuspended in 30 μL 2:1 chloroform:methanol (v/v) and spotted on a silica gel-coated thin-layer chromatography aluminum sheet (Sigma-Aldrich #1055540001). The sheet was developed in an 85:15:5 toluene:chloroform:methanol (v/v/v) solvent to separate TAG from DAG. Images were acquired using Typhoon FLA9500 scanner.

### Generation of fluorescent tag knock-in cell lines

Endogenous tagging of VAPB was performed as described in (Cabukusta et al., 2020). The VAPB/pX330 Cas9 plasmid included the VAPB guide RNA sequence ‘GCCGCTAAGGAACATGGCGA’ (Cho et al., 2022). The VAPB/endomScarlet plasmid was created by swapping the GFP tag in VAPB/endoGFP with an mScarlet tag, and included the left homology arm sequence ‘GCCGTCAGCTCGCCGGGCACCGCGGCCTCGCCCTCGCCCTCCGCCCCTGCGCCTGCACCGCGTAGACCGACCCCCCCCCAGCGCGCCCACCCGGTAGAGGACCCCCGCCCGTGCCCCGACCGGTCCCCGCCTTTTTGTAAAACTTAAAGCGGGCGCAGCATTAACGCTTCCCGCCCCGGTGACCTCTCAGGGGTCTCCCCGCCAAAGGTGCTCCGCCGCTAAGGAACATG’ and the right homology arm sequence ‘GCGAAGGTGGAGCAGGTCCTGAGCCTCGAGCCGCAGCACGAGCTCAAATTCCGAGGTAAGCCCCAGAGGCCGCCACCTTCCTGCCCGCGGCCTCCGCCCCAGTGCTGGAAGGACGGAGCCCGGCGCGGCGGGTGACGTCGGCCCTCGTCCCCACCCCGACGGCGCTGTCGCGGGGGGCGGCGAGGCCGGGCCGGGCCTTGGCGCTCGCCGCGCTCCCCAGAACTGCCCGGAGTGGCCGGGACCCGAGA GGG’.

### GFP immunoprecipitation and western blotting

Endogenously tagged GFP-VAPB MelJuSo cells were seeded in 6 cm dishes. GFP-trap immunoprecipitation was performed as described in (Cabukusta et al., 2020). Western blotting was performed as in (Cabukusta et al., 2020).

### Electron microscopy and ER-LD contact quantification

HeLa cells, transfected with a final concentration of 40 nM siRNA, were cultured in 35 mm dishes. Cells were incubated overnight with 30 μM oleic acid and were fixed for 2 hours at room temperature in 0.1 M Cacodylate buffer containing 1,5% glutaraldehyde. After rinsing thrice with 0.1 M Cacodylate buffer, cells were postfixed with 1% Osmium tetroxide and 1% uranyl acetate. Cells were dehydrated with a series of ethanol, followed by a series of mixtures of ethanol and EPON (LX112, Leadd) and at the end pure EPON. BEEM capsules filed with EPON were placed on the dishes with the open face down. After EPON polymerization at 45°C the first night and 70°C the second night, the BEEM capsules were snapped off. Ultrathin sections 100 nm were made parallel to the surface of the BEEM capsules containing the cultured cells using a Leica EM Ultracut 6. The sections were contrasted with 7% uranylacetate for 20 minutes and Reynolds lead citrate for 10 minutes, and examined at 120 keV with a Tecnai Twin transmission electron microscope (Thermo Fisher, Eindhoven, Netherlands). Overlapping images were automatically collected with a Gatan Oneview camera (Gatan, Pleasonton) at binning 2 at a nominal magnification of 11000 x (1.96 nm pixel size) and stitched together into a composite image as previously described (Faas et al., 2012). Supervised machine learning-assisted analysis was performed using a custom web-based user interface. A subset of LDs was annotated manually in the composite images, and ROI boxes were placed to extract to generate a training set data. A model for the LDs was trained using Keras and TensorFlow, using an adapted version (manuscript in prep.) of a pre-existing convolutional neural network (Xiao et al., 2018). LDs were predicted in all composite TEM images. The ER was annotated manually. The amount of ER-LD contacts was quantified by calculating the percentage of LDs that have contact with the ER annotation in a section of an electron microscopy slice. Contact was defined when ER and LD annotations overlapped. For the quantification of ER-LD contact length, masks of LDs and ER were created in ImageJ (Fig. S7). The border of the ER mask was generated and overlayed with the LD mask using Image Calculator. The length of the overlap of the masks was quantified.

For 3D reconstructions of ER-LD contacts, fixed cells were incubated for 1 hour on ice in Cacodylate buffer containing 1% Osmium tetroxide and subsequently in water containing 1% uranyl acetate. The cells were then dehydrated through a series of ethanol and embedded in EPON. The flat embedded cells were sectioned with an ultramicrotome using a diamond knife at a nominal section thickness of 100 nm. The sections were transferred to a formvar and carbon coated copper single slot grid and stained for 20 minutes with 3% uranyl acetate in water and for 10 minutes with lead citrate according to Reynolds. Electron microscopy images of (∼10) serial sections were recorded at 6500x (3.4 nm/pixel) using a Tecnai 12 TEM (FEI, currently Thermo Fisher Scientific) equipped with an EAGLE 4x4K digital camera. The serial sections were aligned using Amira software. Manual annotation of subcellular structures and creation of images and movies were also performed in Amira.

### TurboID proximity labeling and proteomics

HeLa cells were seeded in 6 cm culture dishes and transfected with the TurboID constructs. The day after transfections, cells were incubated with culture medium containing 50 μM Biotin for 30 minutes. Samples were processed as described in (Cabukusta et al., 2020). Protein samples were stained with silver using SilverQuest Silver Staining Kit (LC6070) for visualization.

For proteomics analysis, TurboID experiments were performed thrice and protein samples were run for 3 cm on a 4%–12% PAGE gel (NuPAGE Bis-Tris Precast Gel, Life Technologies). Gel bands were reduced with 10 mM dithiothreitol, alkylated with 50 mM iodoacetamide. In-gel trypsin digestion was performed using a Proteineer DP digestion robot (Bruker). Tryptic peptides were extracted from the gel slices using 50/50/0.1 water/acetonitrile/formic acid (v/v/v), after which samples were lyophilized. The samples were dissolved in water/formic acid (100/0.1 v/v) and analyzed by online C18 nanoHPLC MS/MS with a system consisting of an Ultimate3000nano gradient HPLC system (Thermo, Bremen, Germany) and an Exploris480 mass spectrometer (Thermo). Peptides were injected onto a cartridge precolumn (300 μm × 5 mm, C18 PepMap, 5 um, 100 A, and eluted via a homemade analytical nano-HPLC column (30 cm × 75 μm; Reprosil-Pur C18-AQ 1.9 um, 120 A (Dr. Maisch, Ammerbuch, Germany). The gradient was run from 2% to 40% solvent B (20/80/0.1 water/acetonitrile/formic acid (FA) v/v) in 30 minutes. The nano-HPLC column was drawn to a tip of ∼10 μm and acted as the electrospray needle of the MS source. The mass spectrometer was operated in data-dependent Top 20 MS/MS mode, with a HCD collision energy at 30% and recording of the MS2 spectrum in the orbitrap, with a quadrupole isolation width of 1.2 Da. In the master scan (MS1) the resolution was 120,000, the scan range 400-1500, at standard AGC target @maximum fill time of 50 ms. A lock mass correction on the background ion m/z=445.12003 was used. Precursors were dynamically excluded after n=1 with an exclusion duration of 10 seconds, and with a precursor range of 20 ppm. Charge states 2-5 were included. For MS2 the first mass was set to 110 Da, and the MS2 scan resolution was 30,000 at an AGC target of 100% @maximum fill time of 60 ms.

In a post-analysis process, Proteome Discoverer 2.5 (Thermo) was used with 10 ppm and 20 ppm deviation for precursor and fragment mass, respectively and trypsin as enzyme was specified. Oxidation on Met and acetylation on N-term were set as a common modification; carbamidomethyl on Cys was set as a fixed modification. An FDR of 1% was set.

The raw data is uploaded to PRIDE under submission number PXD073300.

### Statistical analysis

Statistical analysis was performed with Student’s t test, Mann-Whitney U test or paired Student’s t test using R Software or GraphPad Prism. p values are indicated by asterisks: ∗ is p < 0.05, ∗∗ is p < 0.01, ∗∗∗ is p < 0.001, ∗∗∗∗ is p < 0.0001.

### Reagents

| siRNAs |  |  |
| --- | --- | --- |
| Name | Company | Catalog # |
| siVAPA | Dharmacon | MQ-021382-01-0002 |
| siVAPB | Dharmacon | MQ-017795-00-0002 |
| MOSPD1 | Dharmacon | MQ-020537-00-0002 |
| MOSPD2 | Dharmacon | MQ-017039-01-0002 |
| MOSPD3 | Dharmacon | MQ-016132-02-0002 |
| siAKAP11 | Dharmacon | M-009277-01-0005<br>MQ-009277-01-0002 |
| siOSBPL2 | Dharmacon | M-010361-00-0005 |
| siVPS13A | Dharmacon | M-012878-01-0005 |
| siRAI14 | Dharmacon | M-013919-00-0005 |
| siCOPA | Dharmacon | M-011835-01-0005 |
| siPITPNM1 | Dharmacon | M-019888-00-0005 |
| siPITPNM2 | Dharmacon | M-014106-01-0005 |
| siVPS13C | Dharmacon | M-020641-01-0005 |
| siRAB3GAP1 | Dharmacon | M-032584-01-0005 |
| siLSG1 | Dharmacon | M-031865-00-0005 |
| siWDR44 | Dharmacon | M-018913-00-0005 |
| siINF2 | Dharmacon | M-014097-01-0005 |
| siOSBPL1A | Dharmacon | MQ-008350-02-0002 |

| Antibodies |  |  |
| --- | --- | --- |
| Name | Company | Catalog # |
| ATGL | Abcam | AB220738 |
| ATGL | Proteintech | 55190-1-AP |
| Perilipin-2 | Proteintech | 15294-1-AP |
| VAPB | Proteintech | 14477-1-AP |
| VPS13A | Proteintech | 28618-1-AP |
| ORP1L | In-house | (Johansson et al., 2003) |
| AKAP11 | Proteintech | 30552-1-AP |
| Vinculin | Sigma-Aldrich | V9131 |
| β-Actin | Sigma-Aldrich | A5441 |
| GFP | Chromotek | pabg1 |
| Scarlet | Chromotek | 6g6 |
| Anti-rabbit, Alexa Fluor 488 | Thermo Fisher | A-21206 |
| Anti-rabbit, Alexa Fluor 568 | Thermo Fisher | A-11036 |
| Anti-rabbit, HRP | Thermo Fisher | G-21234 |
| Anti-mouse, HRP | Thermo Fisher | G-21040 |
| Anti-mouse, IRDye 800CW | LI-COR | 926-32210 |

| Plasmids |  |  |
| --- | --- | --- |
| Name | Source | Description |
| VAPB/pX330 | This study | N/A |
| VAPB/endoGFP | (Jongsma, Neefjes et al., manuscript in prep.) | N/A |
| VAPB/endomScarlet | This study | N/A |
| mRFP-C1 | (Cabukusta et al., 2020) | N/A |
| mRFP-VAPB | This study | N/A |
| mRFP-VAPB** | This study | N/A |
| mRFP-VAPB-TM | This study | N/A |
| 2xHA-TurboID-C1 | (Cremer et al., 2023) | N/A |
| 2xHA-TurboID-VAPB | This study | N/A |
| 2xHA-TurboID-VAPB** | This study | N/A |
| 2xHA-TurboID-VAPB-TM | This study | N/A |

| Chemicals |  |  |
| --- | --- | --- |
| Name | Company | Catalog # |
| DharmaFECT 1 | Dharmacon | T-2001 |
| X-treme GENE HP DNA | Sigma-Aldrich | 6366244001 |
| Nile Red | Santa Cruz | sc-203747 |
| BODIPY | Sigma Aldrich | 790389 |
| LysoTracker Deep Red | Thermo Fisher | L12492 |
| NBD-DAG | Cayman Chemical | 9000341 |
| Oleic acid | Sigma-Aldrich | O3008 |
| NG-497 (ATGLi) | Cayman Chemical | 36886 |
| Lalistat 2 (LALi) | Sigma Aldrich | SML2053 |
| T-863 (DGAT1i) | Cayman Chemical | 25807 |
| PF-06424439 (DGAT2i) | Cayman Chemical | 17680 |
| PKI 14-22 | MedChemExpress | HY-P1291A |
| KT 5720 | Cayman Chemical | 10011011 |
| Forskolin | MedChemExpress | HY-15371 |
| GFP-Trap Agarose | Chromotek | gta |
| Biotin | Sigma-Aldrich | B4639 |
| Pierce High Capacity NeutrAvidin Agarose | Thermo Fisher | 29204 |

| Cell lines |  |  |
| --- | --- | --- |
| Name | Source | Description |
| HeLa | (Cabukusta et al., 2020) | N/A |
| MelJuSo | (Cabukusta et al., 2024) | N/A |
| U2OS | ATCC | HTB-96 |
| Huh-7 | (Harryvan et al., 2025) | N/A |
| Hep-G2 | ATCC | HB-8065 |
| HeLa mScarlet-VAPB | This study | N/A |
| MelJuSo GFP-VAPB | (Jongsma, Neefjes et al., manuscript in prep.) | N/A |

| Software and algorithms |  |  |
| --- | --- | --- |
| Name | Source | Description |
| ImageJ (1.53t) | (Schneider et al., 2012) | https://imagej.net/ij/ |
| R | N/A | https://cran.r-project.org/ |
| GraphPad Prism (10.2.3 (403)) | N/A | https://www.graphpad.com/ |
| FFAT score calculation | (Mikitova and Levine, 2012) | N/A |
| FlowJo (v.10) | N/A | https://www.flowjo.com/ |
| Ilastik (1.4.0) | (Berg et al., 2019) | https://www.ilastik.org/ |
| Keras | (Chollet, 2015) | https://keras.io |
| TensorFlow | (Abadi et al., 2015) | http://tensorflow.org |
| MRI Lipid Droplets Tool | N/A | https://github.com/MontpellierRessourcesImagerie/imagej_macros_and_scripts/wiki/Lipid-Droplets-Tool |
| Nearest neighbor distance | N/A | https://www.cavs.msstate.edu/icme/code/MatLab/NND.php |
| TrackMate (7.10.2) | (Ershov et al., 2022; Tinevez et al., 2017) | https://imagej.net/plugins/trackmate/ |
| Shotgun Lipidomics Assistant | (Su et al., 2021) | https://www.uclalipidomics.net/sla.html |
| Amira (2024.2) | N/A | https://www.thermofisher.com/nl/en/home/electron-microscopy/products/software-em-3d-vis/amira-software.html |
| Proteome Discoverer (2.5) | N/A | https://www.thermofisher.com/nl/en/home/industrial/mass-spectrometry/liquid-chromatography-mass-spectrometry-lc-ms/lc-ms-software/multi-omics-data-analysis/proteome-discoverer-software.html |

## Acknowledgements

We thank the members of the Neefjes lab for their valuable input. We thank Lukas J.A.C. Hawinkels for the Huh-7 and Hep-G2 cell lines. We thank Evelyne Steenvoorden for technical assistance.

## Funding

This research is supported by the Netherlands ALS Foundation (“*Stichting ALS Nederland*”, AV20240007) awarded to Birol Cabukusta, and by the Spinoza award of the Dutch Research Council (“*Nederlandse Organisatie voor Wetenschappelijk Onderzoek*”; NWO 00897590) awarded to Jacques Neefjes. Jacques Neefjes is an Oncode Institute investigator.

## Author Contributions

Conceptualization of the study: SBP, MS, MG, JN and BC; methodology, validation, statistical analysis, visualization: SBP and BC; investigation: SBP, MS, LLJJ, SR, AHdR, AWMdJ, EB and BC; access to facilities: PAvV, RIK and MG; supervision of the study: MG, JN, and BC; project coordination and funding acquisition: BC and JN. SBP and BC wrote the original manuscript with critical input from MG, JN and other authors.

## Competing Interests

Authors declare no competing interest.

---

**Supplementary figure 1.**
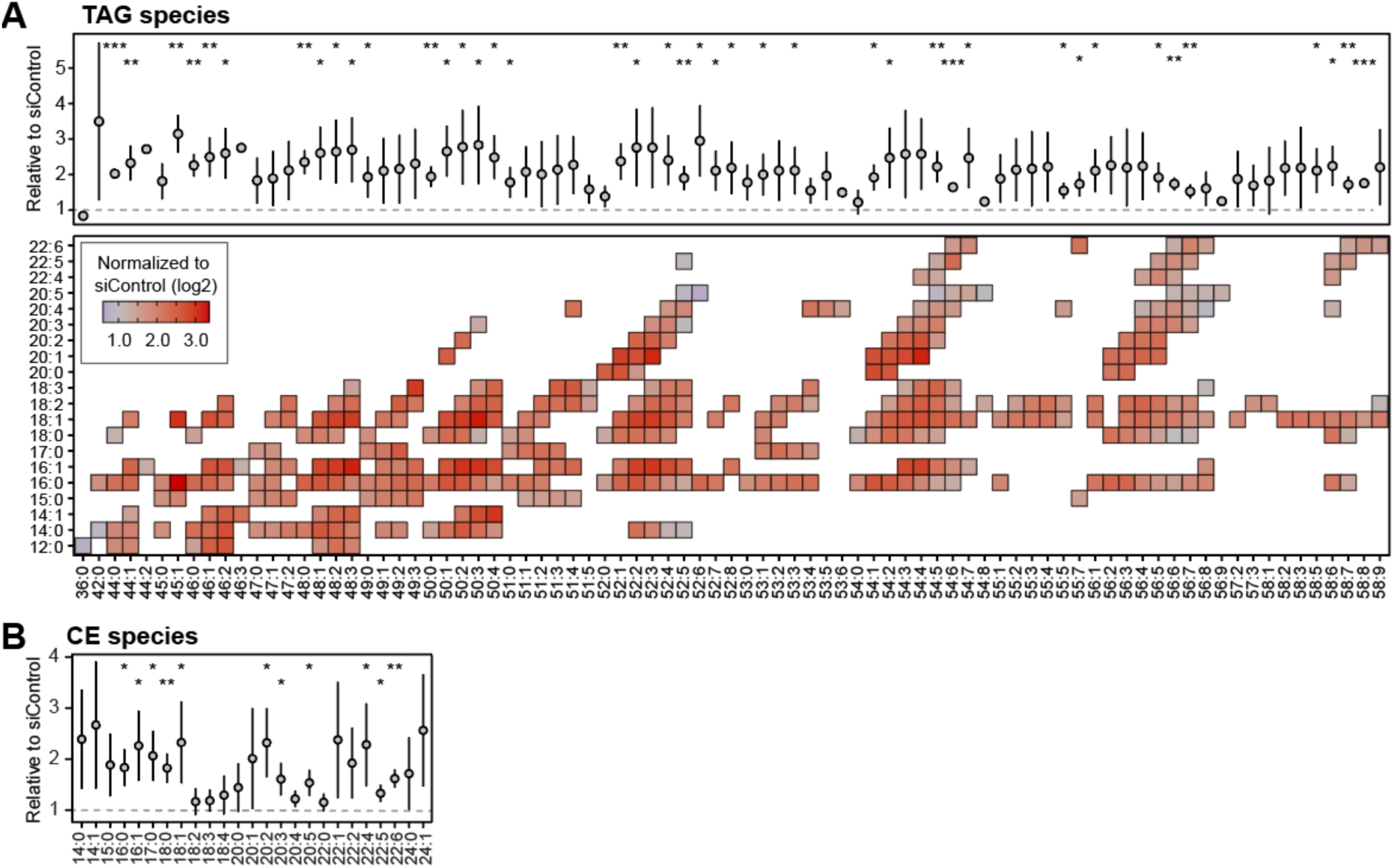
Nearly all TAG and CE species are increased in VAPB depleted cells. **(A and B)** Lipidomic analysis of intracellular TAG species (A) and CE species (B) of siVAPB-treated cells. Data is relative to siControl. Results are mean ± SD of three independent experiments.

**Supplementary figure 2.**
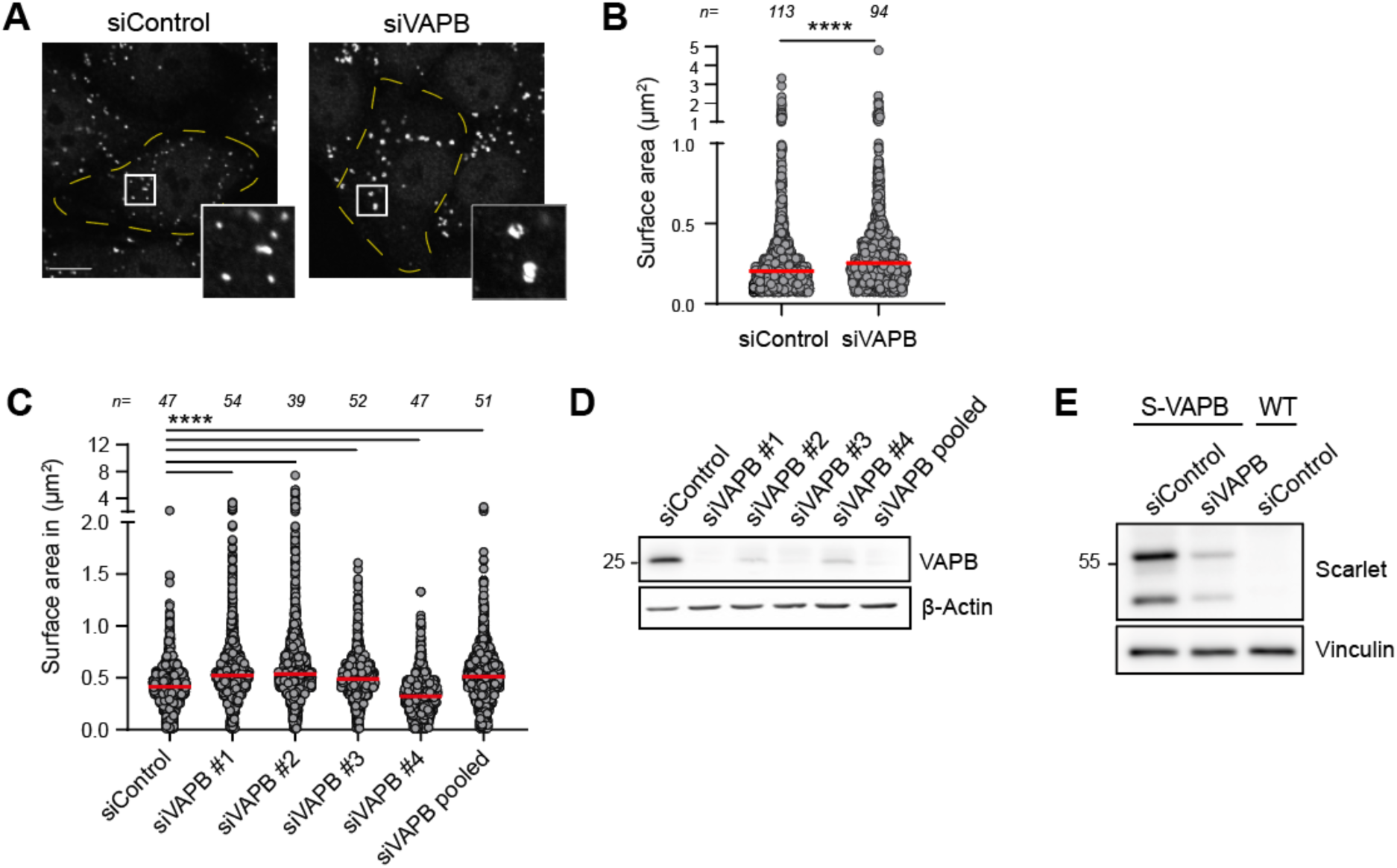
siRNAs targeting VAPB are confirmed to deplete VAPB and increase LD size. **(A)** Representative fluorescence images of siControl- and siVAPB-treated cells stained against PLIN2. Scale bar: 10 µm. **(B)** Quantification of LD surface area corresponding to images of panel A. Red bars represent the median of one experiment; n is the number of analyzed cells. **(C)** Quantification of LD surface area of cells treated with single and pooled VAPB-targeted siRNAs. Red bars represent the median of one experiment; n is the number of analyzed cells. **(D)** Western blot of cells treated with single and pooled VAPB-targeted siRNAs showing VAPB depletion efficiency. **(E)** Western blot validation of endogenous tagging of VAPB with mScarlet (S-VAPB) by siControl and siVAPB treatment.

**Supplementary figure 3.**
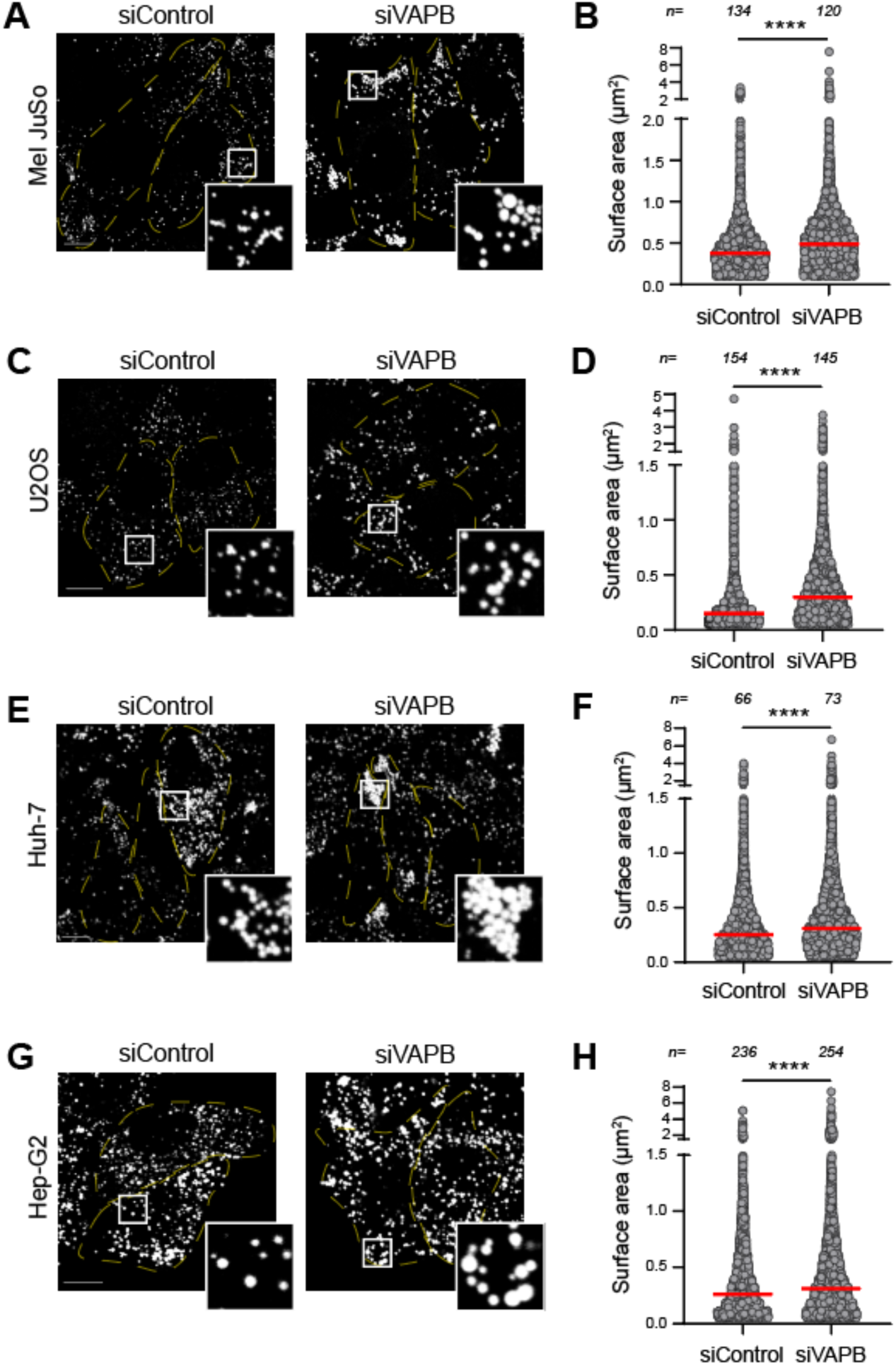
VAPB regulation of LD size is observed in multiple cell lines. **(A, C, E and G)** Representative fluorescence images of siControl- and siVAPB-treated MelJuSo (A), U2OS (C), Huh7 (E) and HepG2 (G) cells stained with BODIPY to visualize LDs. Scale bar: 10 µm. **(B, D, F and H)** Quantification LD surface area corresponding to images of panel A, C, E and G, respectively. Red bars represent the median of three independent experiments; n is the number of analyzed cells.

**Supplementary figure 4.**
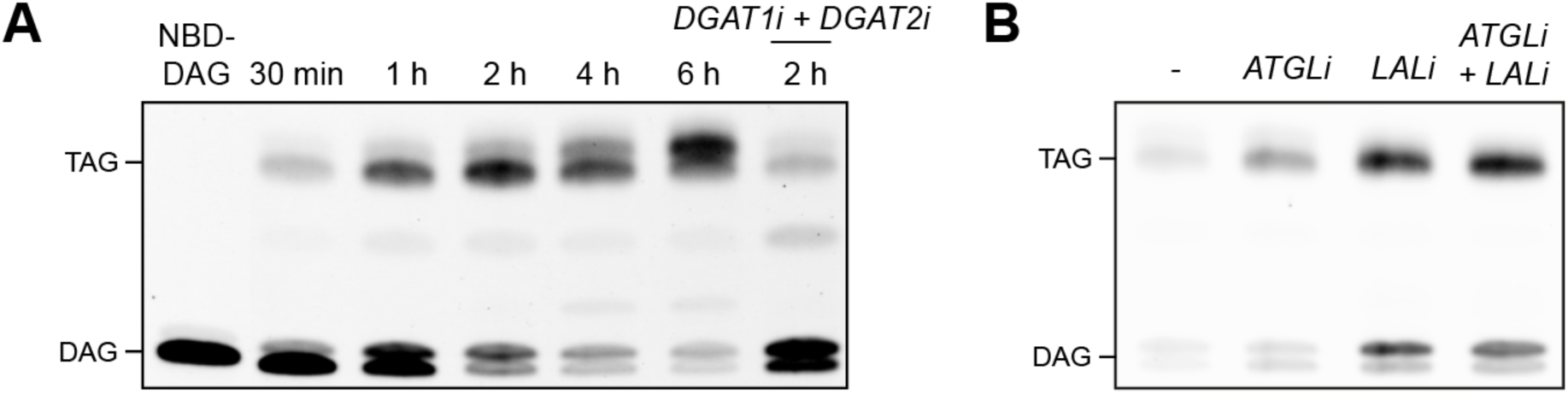
NBD-TAG is synthesized from NBD-DAG and its degradation is inhibited by ATGLi and LALi. **(A)** Thin-layer chromatography image of metabolic chasing experiment where cells were incubated with NBD-DAG for various lengths of time, as indicated. Addition of TAG synthesis inhibitors (DGAT1i+DGAT2i) confirms the identity of the NBD-TAG band. **(B)** Thin-layer chromatography image of cells incubated with NBD-DAG and ATGLi or LALi for 1 hour, depicting accumulation of NBD-TAG.

**Supplementary figure 5.**
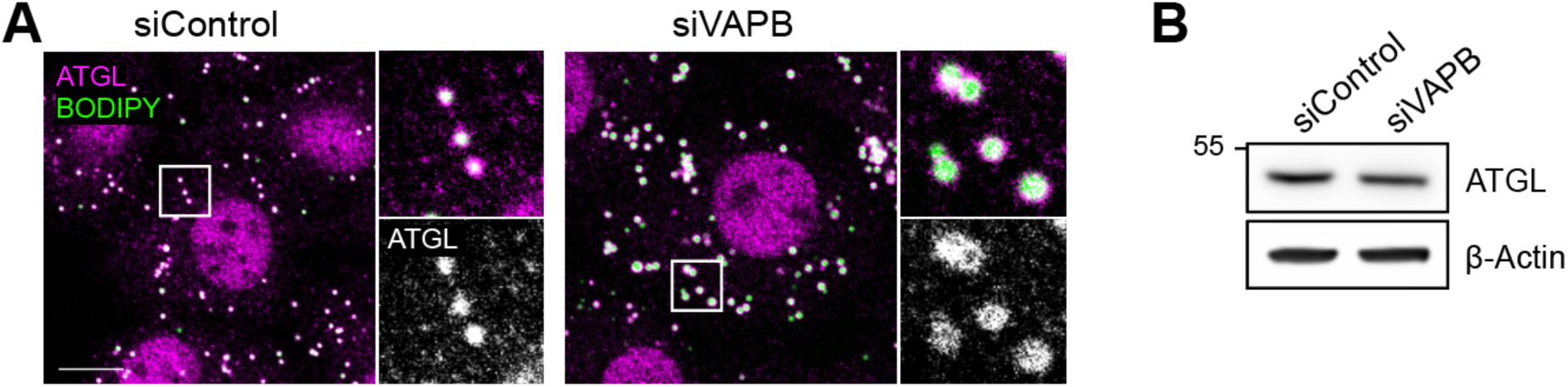
VAPB depletion does not alter ATGL localization or protein levels. **(A)** Immunofluorescence images of siControl- and siVAPB-treated cells stained against ATGL and stained with BODIPY to visualize LDs. Scale bar: 10 µm. **(B)** Western blot of ATGL levels from siControl- and siVAPB-treated cells. Scale bar: 10 µm.

**Supplementary figure 6.**
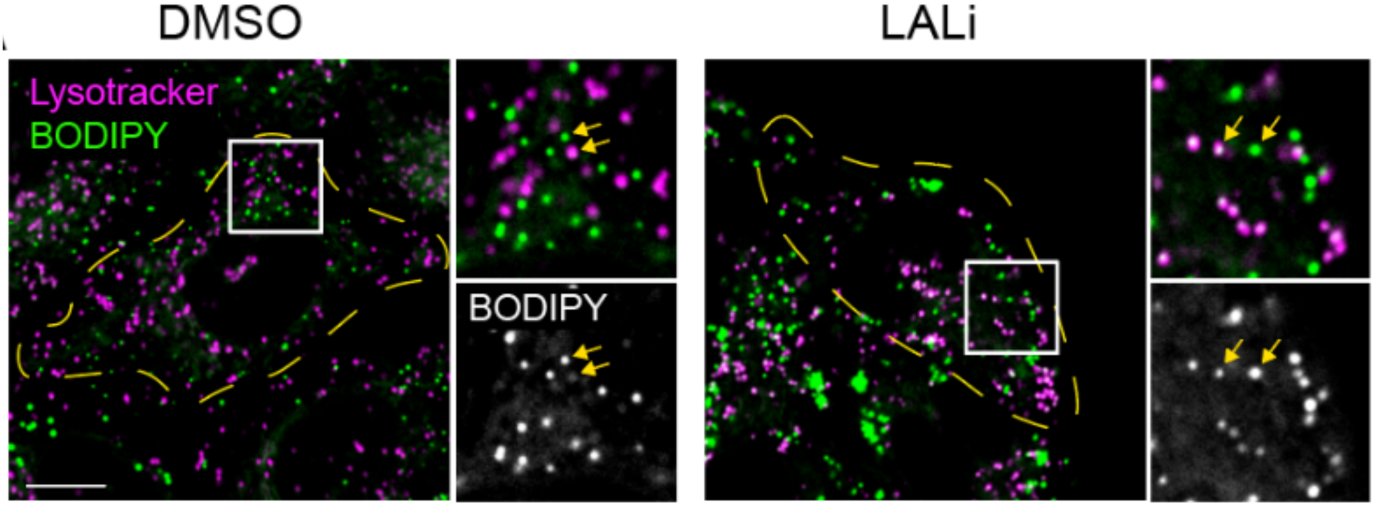
LD retention in lysosomes after LALi treatment. Fluorescence images of cells incubated with DMSO or LALi and stained with BODIPY and lysotracker to label LDs and lysosomes, respectively. Note the accumulation of BODIPY in lysosomes in LALi-treated cells. Scale bar: 10 µm.

**Supplementary figure 7.**
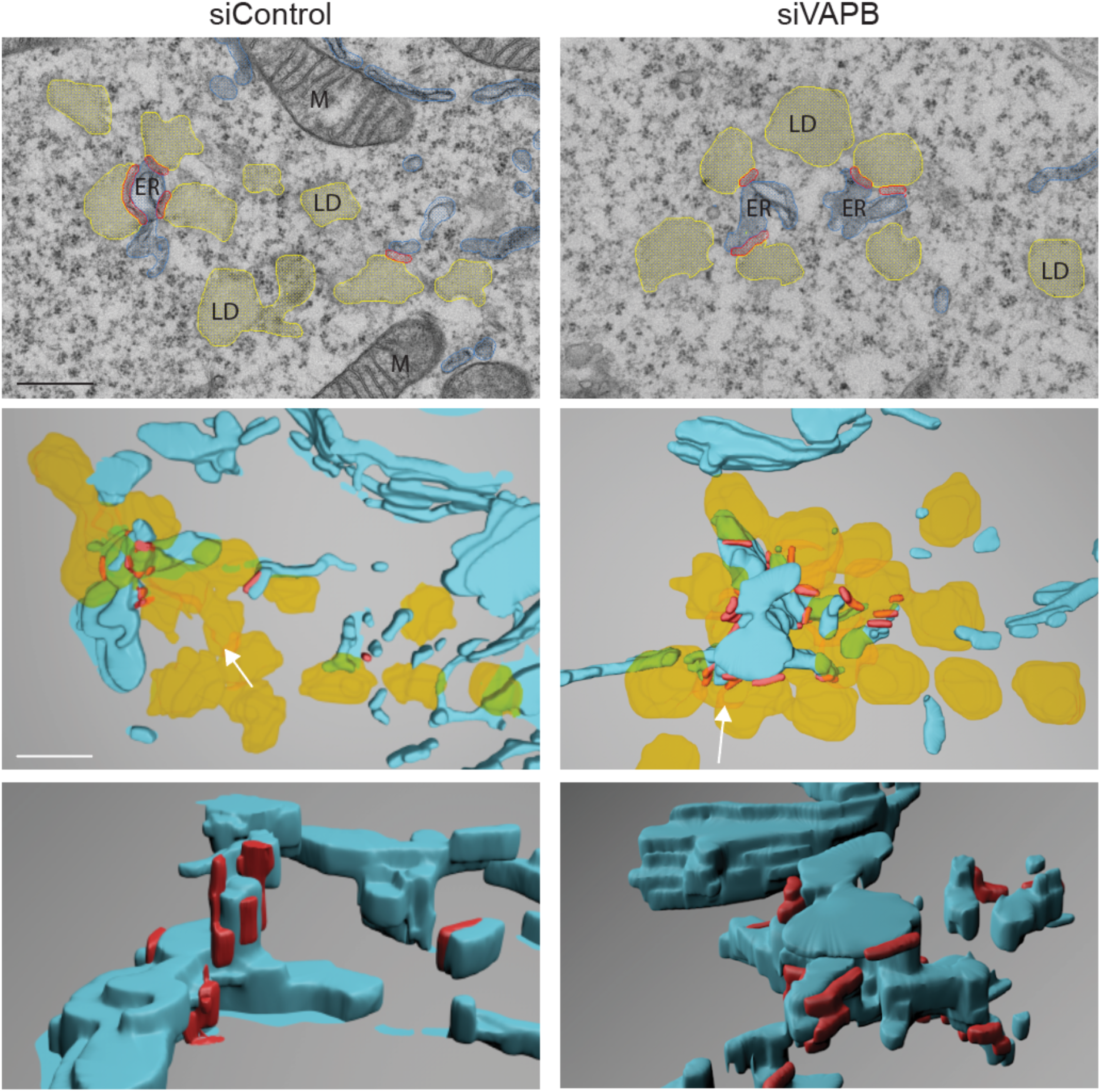
3D reconstructions of ER-LD contacts. Electron microscopy images and corresponding 3D reconstructions from siControl and siVAPB cells. ER, LD and ER-LD contacts are annotated in blue, yellow and red, respectively. M: mitochondria. Scale bar: 500 nm.

**Supplementary figure 8.**
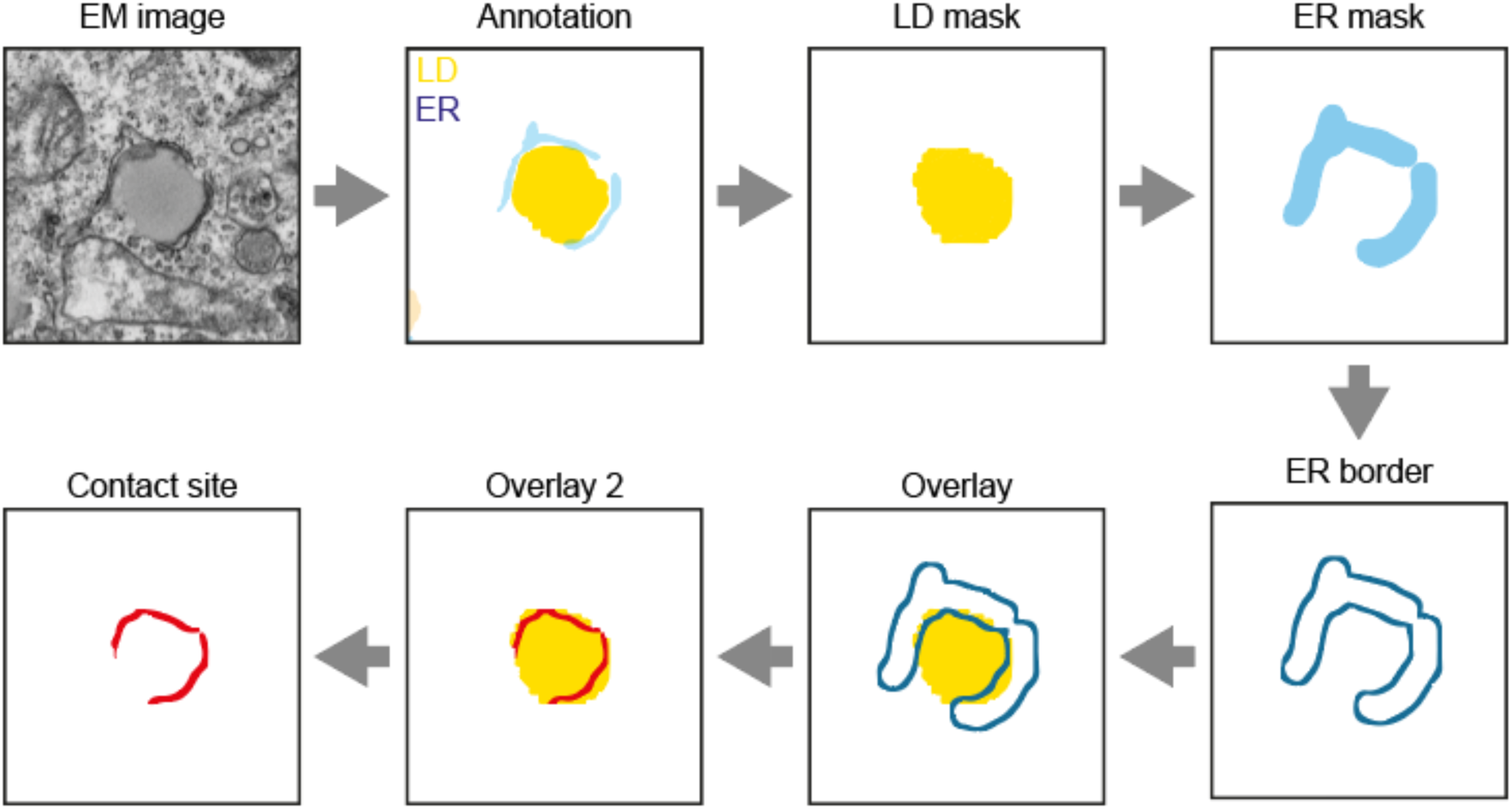
Workflow of ER-LD contact site quantification. The length of ER-LD contacts was quantified by annotating the ER and LDs manually and automatically, respectively. An LD mask was generated based on the annotation, and an ER mask was created by eroding (i.e. enlarging) the ER annotation. Next, the border of the ER mask was created and overlayed with the LD mask. The overlap of the ER border on the LD mask was defined as a contact site and quantified.

**Supplementary figure 9.**
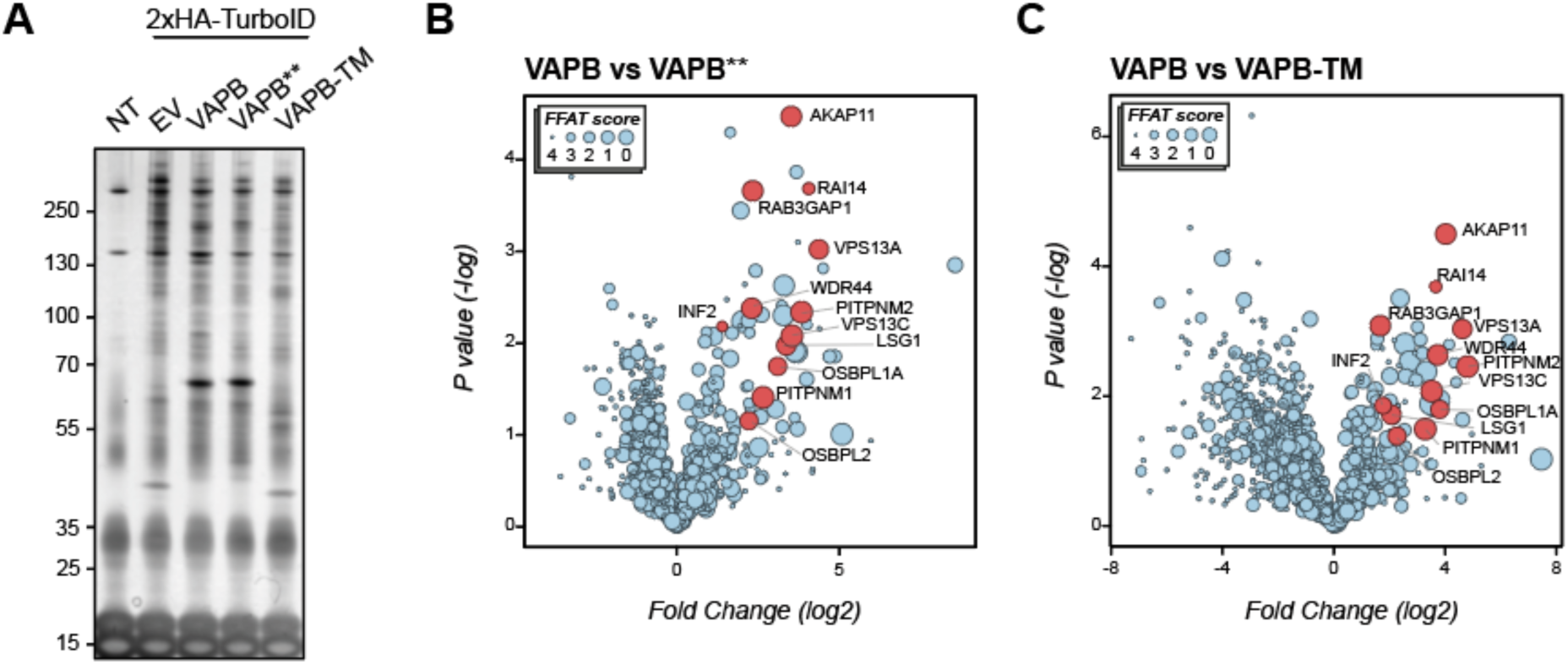
TurboID identifies proteins enriched in VAPB interactome. **(A)** Silver staining of non-transfected (NT), 2xHA-TurboID-empty vector (EV), and 2xHA-TurboID-tagged VAPB, VAPB** and VAPB-TM streptavidin pull-down samples. **(B and C)** Volcano plots of proteins enriched in wild-type VAPB in comparison to VAPB** (B) and VAPB-TM (C). Proteins chosen for validation are depicted in red.

**Supplementary table 1.** Lipidomic analysis of HeLa cells depleted of individual VAP proteins, corresponding to Figure 1B. CE: cholesterol ester, CER: ceramide, DAG: diacylglycerol, DCER: dihydroceramide, FFA: free fatty acid, HCER: hexosylceramide, LCER: lactosylceramide, LPC: lysophosphatidylcholine, LPE: lysophosphatidylethanolamine, PC: phosphatidylcholine, PE: phosphatidylethanolamine, SM: sphingomyelin, TAG: triacylglycerol.

**Supplementary movie 1.**
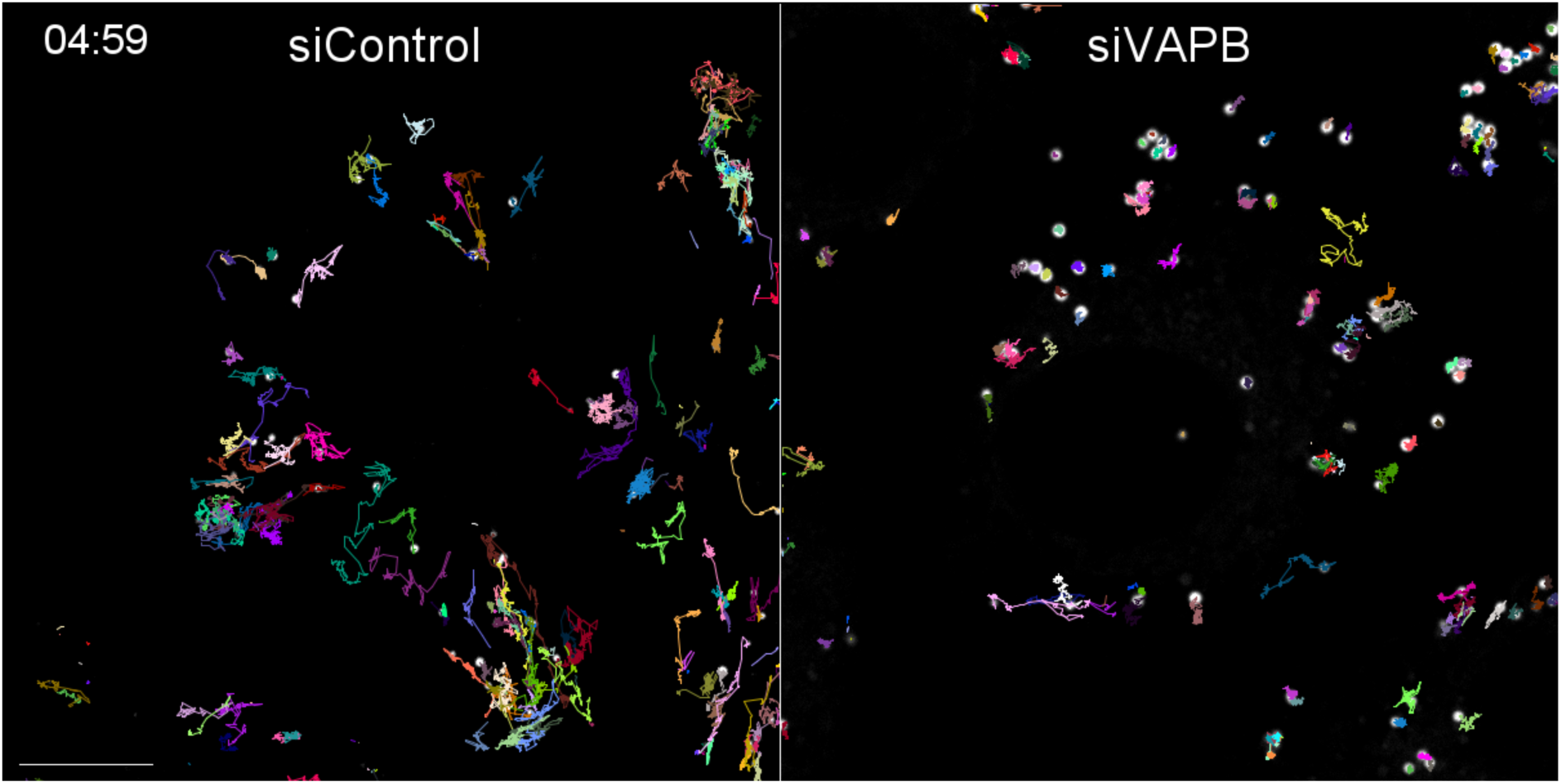
VAPB depletion impairs LD motility. TrackMate tracks of LD movement in BODIPY-stained control- and VAPB-depleted cells. Scale bar: 10 µm.

**Supplementary movie 2.**
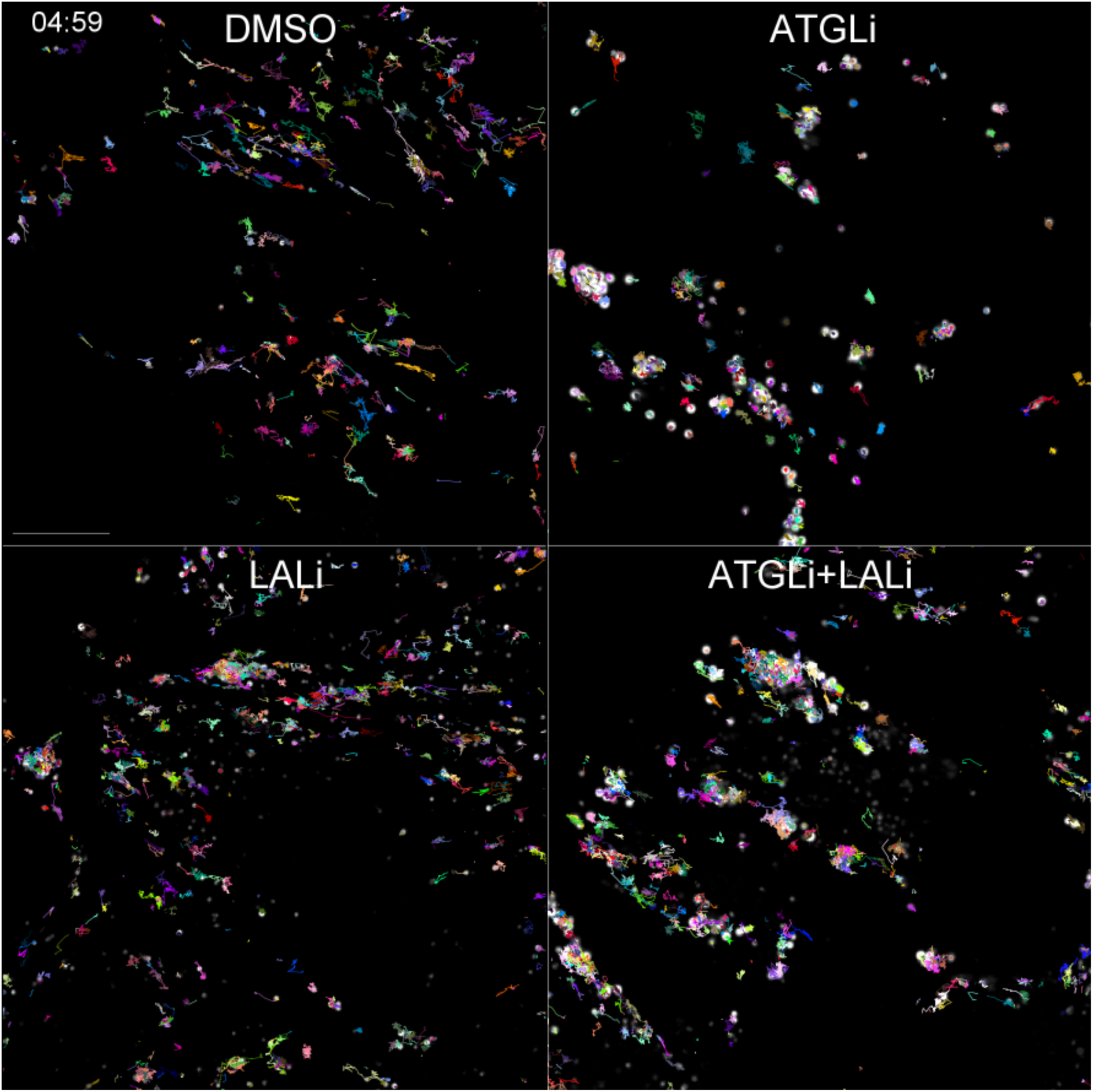
Inhibition of ATGL decreases LD motility. TrackMate tracks of LD movement in cells treated with ATGLi or LALi, stained with BODIPY to visualize LDs. Scale bar: 10 µm.

**Supplementary movie 3.**
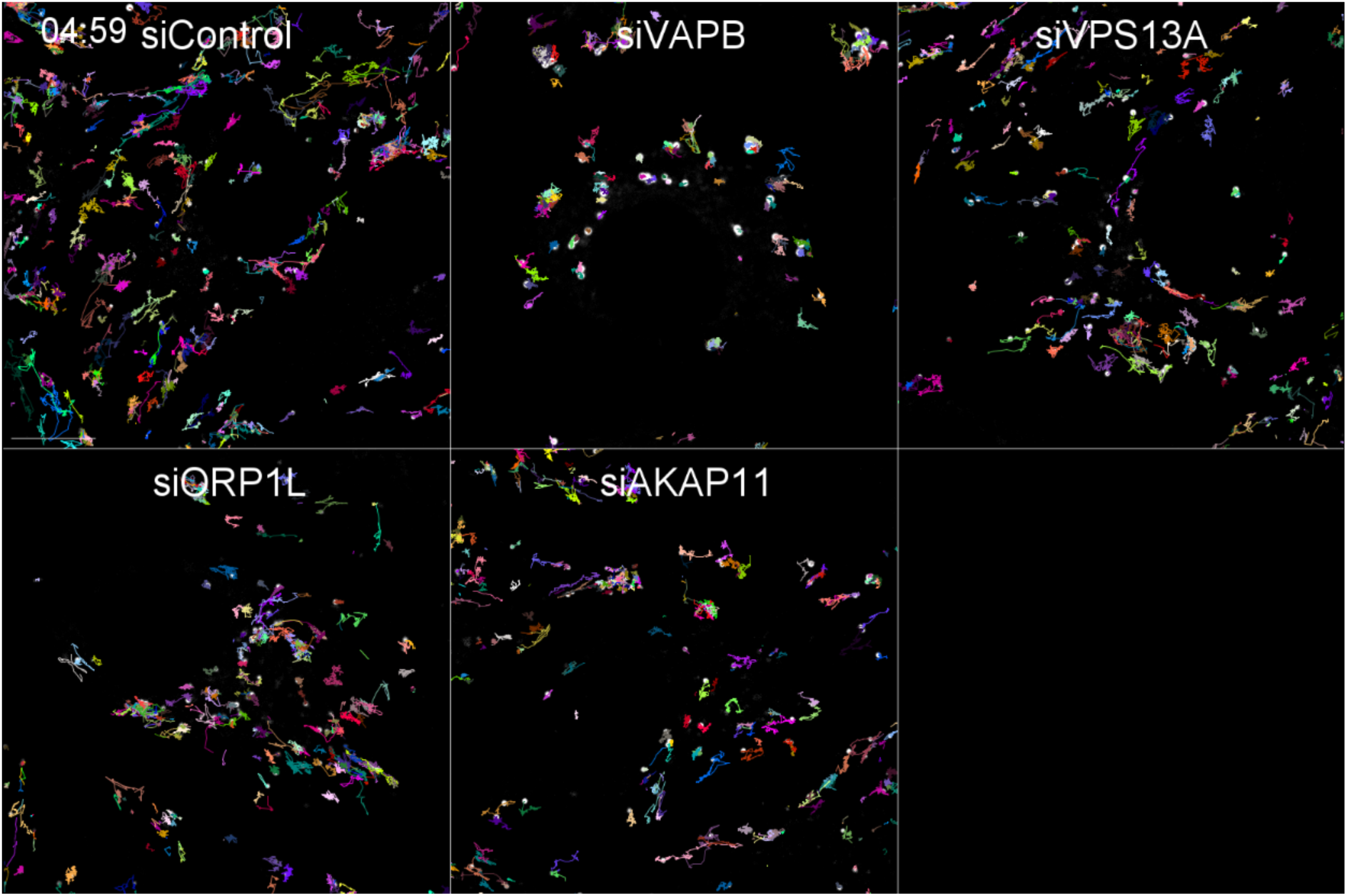
LD motility is affected by VAPB interaction partners. TrackMate tracks of LD movement in cells depleted of VAPB, VPS13A, ORP1L or AKAP11 stained with BODIPY. Scale bar: 10 µm.

## Notes

### Competing Interest Statement

The authors have declared no competing interest.

